# Structure of the *Chimalliviridae* bacteriophage Goslar reveals host recognition and infection machinery

**DOI:** 10.64898/2026.04.21.720040

**Authors:** Dwaipayan Basu, Yajie Gu, Ying-Xing Li, Taylor Forman, Majid Ghassemian, Joe Pogliano, Kevin D. Corbett

**Affiliations:** Department of Chemistry & Biochemistry, University of California San Diego, La Jolla CA 92093 USA; Department of Cellular & Molecular Medicine, University of California San Diego, La Jolla CA 92093 USA; Department of Molecular Biology, University of California San Diego, La Jolla CA 92093 USA

## Abstract

Large-genome bacteriophages (phages) of the *Chimalliviridae* family possess a distinctive life cycle, building compartments inside infected bacterial cells that broadly protect their genomes from host defenses. Here, we use cryoelectron microscopy to determine a high-resolution structure of the *E. coli*-infecting *Chimalliviridae* phage Goslar, generating a composite model with 2,888 protein chains from 28 different structural proteins, totaling over 1.4 million amino acid residues. Our structure reveals the architecture of the Goslar capsid, portal, tail, and baseplate, highlighting structural similarities and differences with other tailed phages. Combining our structural data with quantitative mass spectrometry of Goslar virions, we identify several high-copy virion-associated proteins that likely play key roles when injected into host cells upon infection. We also identify a *Chimalliviridae*-specific family of proteins that form long, flexible filaments anchored at the phage baseplate and which incorporate carbohydrate-binding domains, suggesting a key role in host recognition.

## INTRODUCTION

Bacteriophages (phages) are the most abundant organisms on Earth, outnumbering their bacterial prey by an order of magnitude^1,2^. Phages show huge diversity in their genome sizes — ranging from 3-5 kb for *Microvidiae*^3^ to over 800 kb for so-called jumbo phages^4–6^ — as well as in their physical morphology and overall life cycle. The most well-studied phages possess relatively small genomes, typified by phages M13 (6.4 kb), lambda (48.5 kb), and T4 (168.9 kb). Historically, jumbo phages (classified as those with genomes over 200 kb) have been understudied because typical methods did not allow for their isolation. Recent studies using deep-sequencing data from environmental samples^4^ and/or new isolation methods^7,8^, however, have revealed that jumbo phages are abundant and highly diverse^9^.

One family of jumbo phages, termed *Chimalliviridae*, has a distinctive life cycle that enables it to evade DNA-targeting host defenses like CRISPR-Cas nucleases and restriction endonucleases^10–15^. Upon initial infection, *Chimalliviridae* phages (here termed chimalliviruses) inject their genome into a membrane-enclosed vesicle termed the EPI (early phage infection) vesicle^12,13,16^. A suite of phage proteins is injected into the host cell alongside the genome, including a multisubunit RNA polymerase (the virion RNA polymerase or vRNAP)^17–19^, which transcribes early genes from within the EPI vesicle^12,13,16^. One early-expressed protein is called chimallin (“shield”), and this protein self-assembles into a closed compartment termed the phage nucleus, into which the phage genome is transferred from the EPI vesicle^11,20,21^. Inside the phage nucleus, the genome is replicated and late genes are transcribed by a second phage-encoded RNA polymerase (the non-virion RNA polymerase or nvRNAP)^11,13,19,22–25^. Capsids are assembled in the host cytoplasm, then migrate to the phage nucleus, where they dock onto the chimallin shell for genome packaging^11,20,26^. Capsids and tails are then assembled within a subcellular space termed the bouquet, followed by host-cell lysis^26–28^.

Many questions remain regarding the complex life cycle of chimalliviruses. Notably, the segregation of the EPI vesicle and phage nucleus from the host cytoplasm means that transcribed mRNAs must be transported out of these compartments to be translated by host ribosomes; the mechanisms of mRNA transport in these phages is largely unknown^12,29^. Similarly, many phage- and host-encoded proteins are specifically transported into the phage nucleus via an import apparatus whose mechanisms and specificity are poorly characterized^30,31^. Further, while low-resolution structural studies of intact chimalliviruses have been performed^32^, their detailed architecture is not well-understood, meaning that host recognition and infection mechanisms of this phage family remain mostly unknown. Finally, the full complement of proteins injected alongside the chimallivirus genome upon initial infection is not known, leaving open the questions of how early genes are transcribed and how their mRNAs are transported out of the EPI vesicle.

Likely because of their broad resistance to the most abundant host defense pathways, combined with their diverse host-recognition mechanisms, many chimalliviruses show extremely broad host range and hold significant promise as antibacterial therapeutics^8,33^. Addressing how these viruses recognize their host and navigate early-infection events could enable rational selection or engineering of these phages for therapies targeted to particular patient isolates.

Here, we use cryoelectron microscopy (cryoEM) to determine a near-complete structure of one chimallivirus, the *E. coli*-infecting phage Goslar, and use mass spectrometry to comprehensively define its protein payload. Our final molecular model comprises 2,888 individual chains from 28 distinct proteins. Our structure, along with a recent structure of the *Pseudomonas*-infecting phage PhiKZ^34^ together define the detailed architecture of chimalliviruses for the first time, revealing broad similarity to other tailed viruses (*Caudoviricetes*) with several distinctive features. Our mass spectrometry survey of virion-associated proteins reveals that each Goslar virion carries hundreds of copies of several structurally similar proteins thought to comprise an internal capsid structure called the “inner body” and play key roles upon injection into the host cell. Further, we identify a chimallivirus-specific protein family characterized by the DUF7941 domain; these proteins likely mediate host recognition by self-assembling into extended flexible filaments radiating up to several microns from the phage baseplate.

## RESULTS

### CryoEM structure determination of phage Goslar

To determine the overall structure of phage Goslar, we prepared high-titer phage lysates (∼10^9^ plaque-forming units (PFU)/mL) by infecting lawns of *E. coli* MC1000 cells on agar plates, collecting the resulting phage in a stabilizing buffer, then concentrating using centrifugal ultrafiltration. Further purification by PEG precipitation and CsCl gradient centrifugation did not result in higher particle density on cryoEM grids, so we used samples prior to these steps for structure determination, and the more highly purified samples for mass spectrometry analysis (see below). We prepared cryoEM samples on graphene-coated lacey carbon grids, and initially collected a 4,224-micrograph dataset on a Titan Krios G3 with Gatan K3 camera in super-resolution mode, with a super-resolution pixel size of 1.3 Å/pixel (Dataset 1). From this dataset, we manually picked 3,033 intact Goslar phage particles and calculated a ∼19 Å resolution asymmetric (C1) map of the full Goslar virion in cryoSPARC (**Figure S1, Table S1**)^35^. In this map, the capsid, portal/neck, tail, and baseplate are clearly visible (**Figure 1A-B**).

**Figure 1.**
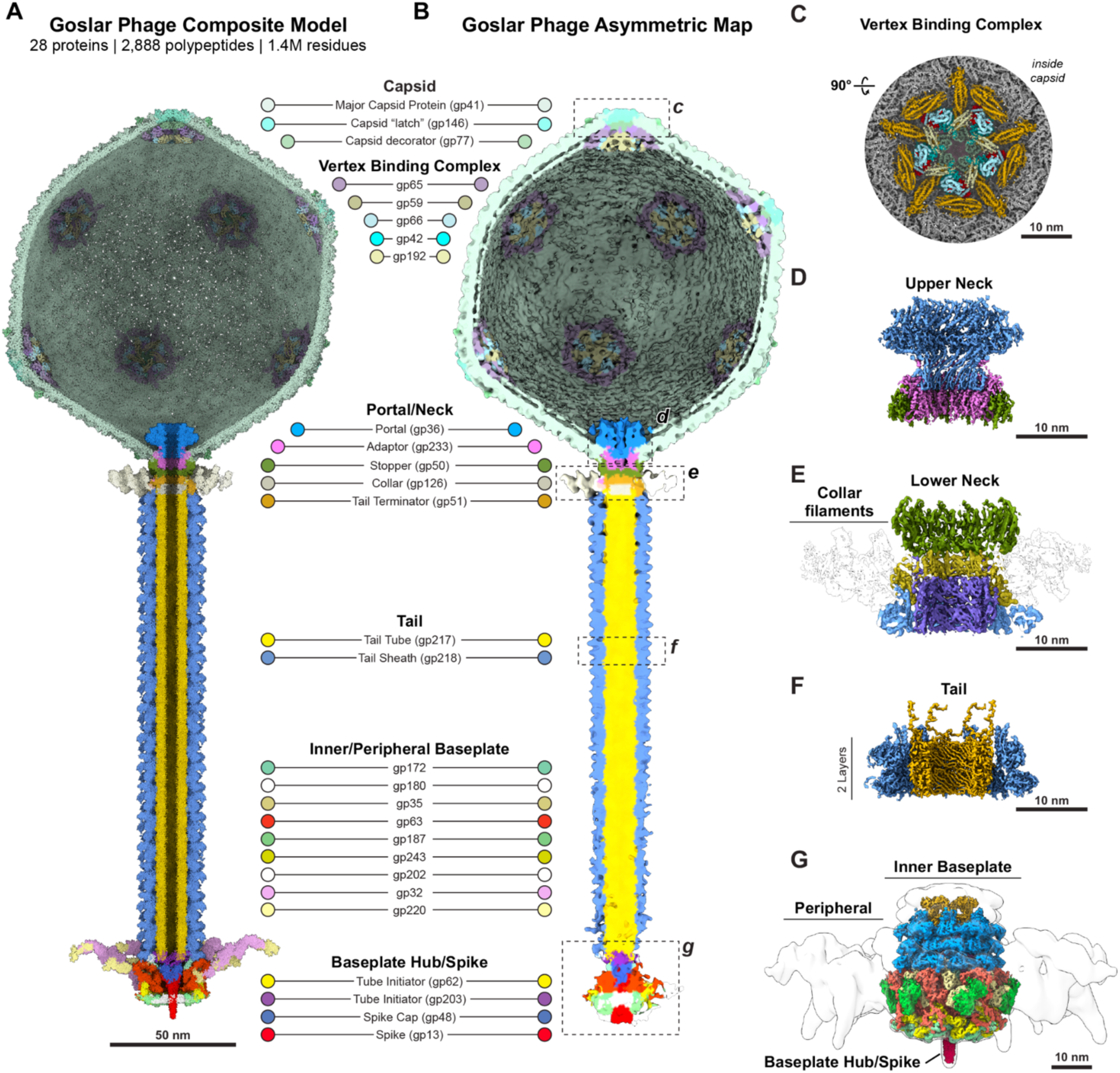
Overview of cryoEM structure determination of the Goslar virion. A. Composite model of the Goslar virion, comprising 2,888 individual polypeptide chains from 28 different proteins (see **Table S3** for copy number of each protein). Proteins are colored according to the key. B. Asymmetric map (∼19 Å resolution) of the Goslar virion, colored by protein as in panel (A). Dotted boxes indicate the positions of detail maps in panels (C) through (G) C. 3.6 Å resolution map of the capsid C5 “penton” vertex, viewed from inside the capsid with the vertex binding complex colored as indicated in panels (A) and (B). See Figure 2 for further details of capsid structure, and Figure 3 for further details of the vertex binding complex. D. 3.8 Å resolution map of the C12-symmetric portal and upper neck, with portal protein (gp36) colored blue, adaptor protein (gp233) pink, and the C12-symmetric N-terminal domains of the stopper protein (gp50). E. 4.2 Å resolution map of the C6-symmetric lower neck. The C6-symmetric C-terminal domains of the stopper protein (gp50) are shown in green, with the tail terminator (gp51) in yellow, the tail tube (gp217) in purple, and the tail sheath (gp218) in blue. The collar and collar filaments (gp126) are shown as a transparent outline. See Figure 4 for further details of the portal/neck and collar. F. 3.4 Å resolution map of the tail (two layers shown), with tail tube protein (gp217) in yellow and tail sheath (gp218) in blue. See Figure 5 for further details of the tail. G. 3.9 Å resolution map of the baseplate, with proteins colored as in panels (A) and (B). The peripheral baseplate is shown as a transparent Gaussian-filtered map. See Figure 6 for further details of the baseplate.

For high-resolution analysis, we separately collected two datasets totaling 13,518 micrographs on the same Titan Krios G3 with Gatan K3 camera in counting mode, with a pixel size of 1.4 Å/pixel (Datasets 2 and 3). The smaller field of view in these micrographs resulted in fewer complete virions being visible, but the resulting reconstructions of individual substructures were higher-quality than those calculated from Dataset 1. Using cryoSPARC^35^, we pursued a multi-pronged strategy for structure determination of intact Goslar virions that leveraged the high symmetry of each substructure. For the capsid, we picked 5,214 intact capsids with attached tails, and generated a 6.3 Å resolution icosahedrally-averaged (I1) map of the entire capsid (**Figure S2-S3, Table S2**). Using this map, we performed symmetry expansion followed by map rotation to align particles to either the three-fold (C3) or five-fold (C5) symmetry axes of the capsid. Using particles aligned on the C3 axis, we generated a 3.6 Å resolution map centered on a hexamer of the major capsid protein (C3 hexon). Using particles aligned on the C5 axis (C5 penton vertex), we first performed 3D classification to identify the single vertex in each capsid that contained the portal. Penton vertex particles without the portal were combined and refined with C5 symmetry to generate a 3.6 Å map centered on a pentamer of the major capsid protein (C5 penton; **Figure 1C**). Portal vertex images were re-centered and refined with C12 symmetry to obtain a 3.8 Å resolution map of the portal and upper neck (**Figure 1D**), and with C6 symmetry to obtain a 4.2 Å resolution map of the lower neck and the top of the tail (**Figure 1E**).

From Dataset 1, we separately picked particles corresponding to empty capsids, and calculated a 5.2 Å resolution icosahedrally-averaged map from 4,535 particles (**Figure S4, Table S2**). We pursued a similar symmetry expansion and 3D classification approach as described above for intact phage particles, but could not identify any portal vertices in the empty capsids. This observation, plus the fact that empty capsids universally lacked tails, suggests that these particles represent capsids that self-assemble into a closed icosahedrally-symmetric structure without a portal.

Next, to determine the structure of the phage tail, we used the cryoSPARC filament tracer to pick 47,640 tail-segment particles from Datasets 2 and 3, then refined with C6 plus helical symmetry (twist 21.5°, rise 39.2 Å) to obtain a 3.4 Å resolution map (**Figure S5, Table S2, Figure 1F**). We separately masked and re-refined the inner tail tube and outer tail sheath substructures, resulting in a 3.2 Å tail tube map and a 3.4 Å tail sheath map (**Figure S5, Table S2**).

Finally, to determine the structure of the baseplate, we used template picking to obtain 5,539 baseplate particles from Datasets 2 and 3, then refined with C6 symmetry to obtain a 3.9 Å map (**Figure S6, Table S2, Figure 1G**). In this map, the “hub” and inner baseplate structures can be confidently assigned to individual proteins; a much larger peripheral baseplate structure that includes the tail fibers could be observed in the map, but was not well defined. To better define this region, we performed symmetry expansion and local refinement using a mask covering 1/6th of the peripheral baseplate region. This strategy resulted in a ∼8 Å resolution map, into which we could tentatively assign the density of the “spokes” connecting the inner and peripheral baseplate structures (see below). We were unable to assign the density of the peripheral baseplate ring or the tail fibers.

To build atomic models into each high-resolution map, we leveraged a combination of sequence homology to known phage structural proteins, sequence fragments assigned by ModelAngelo^36^, 3D models and predicted protein-protein interactions from AlphaFold 3 (ref. ^37^), and mass spectrometry of purified Goslar virions (see below) to identify and build models for each protein visible in the capsid, portal, neck, tail, and baseplate maps. The final models comprise a total of 2,888 separate protein chains from 28 different structural proteins, totaling over 1.4 million amino acid residues.

A primary motivation for our structure determination effort was to visualize the so-called “inner body”, a large structure previously observed in low-resolution cryoEM studies of the *Pseudomonas*-infecting chimallivirus PhiKZ that forms a pillar-like assembly spanning the inside of the filled capsid, positioned diagonally with respect to the portal/tail axis^38,39^. The inner body has been thought to comprise a set of structurally-related proteins that are abundant in the virion^40–42^, but this structure was not resolved in a recent capsid structure of PhiKZ; potentially due to the use of icosahedral symmetry averaging in that work^41^. A more recent study that included a ∼10 Å resolution asymmetric cryoEM structure of the intact PhiKZ virion also did not observe density for the inner body^34^. We did not observe density for the inner body in our primary micrographs of Goslar (**Figure S1A**), or in our asymmetric (C1) map of the Goslar virion (**Figure 1B**), in which particles were unambiguously aligned with respect to both the capsid and portal/neck/tail substructures. To account for potential ambiguous alignment of the inner body in different particles, we performed C5 symmetry expansion of capsid particle images along the portal vertex and performed 3D classification (not shown); this effort also did not reveal density for the inner body. The lack of density for the inner body, combined with our inability to observe this structure in primary micrographs, suggests either that Goslar does not possess a highly-ordered inner body or that our imaging parameters did not allow for visualization of this structure. The Goslar capsid does contain several high-copy virion-associated proteins homologous to proteins from PhiKZ that have been thought to comprise the inner body; these are discussed more below.

### Capsid architecture

The Goslar capsid has a diameter of ∼147 nm (**Figure S4H**) and adopts an overall icosahedral (I1) symmetry (**Figure 2A**), with the exception that at one of the fourteen vertices, a pentamer of the major capsid protein (MCP; gp41) is replaced by the portal (described below). Each of the 60 asymmetric units (**Figure 2B**) is composed of 27 copies of the MCP (triangulation number (T)=27, h=3, k=3). The total MCP copy number in the capsid is therefore 1,615 (27 x 60 (I1 symmetry) = 1,620 - 5 (portal vertex) = 1,615). Each asymmetric unit contains four complete MCP hexamers (hereafter referred to as “hexons”); one-third of a fifth hexon; and one fifth of an MCP pentamer (hereafter referred to as a “penton”) (**Figure 2C**). Our final models of the MCP in the penton and hexon contain 576 and 583 of the total 740 residues, respectively. No density for the N-terminal 157 residues of the MCP were observed in either structure, suggesting that the Goslar MCP is subject to proteolytic processing like the MCP of PhiKZ^40,41^. Each asymmetric unit also contains one copy each of two proteins that decorate the outer capsid surface: the capsid “latch” protein gp146 binds and stabilizes the MCP penton, and the capsid decorator protein gp77 is a small globular protein bound to the outer surface between two adjacent MCP protomers (**Figure 2C**). We clearly observe one copy of gp77 per asymmetric unit bound to hexon 1, but at least two additional copies per asymmetric unit are also visible at contour levels too low to build (not shown). Finally, the asymmetric unit contains five additional proteins (gp42, gp59, gp65, gp66 and gp192) that form a “vertex binding complex” (VBC) on the inner surface of the capsid (**Figure 2C**; described in more detail next section).

**Figure 2.**
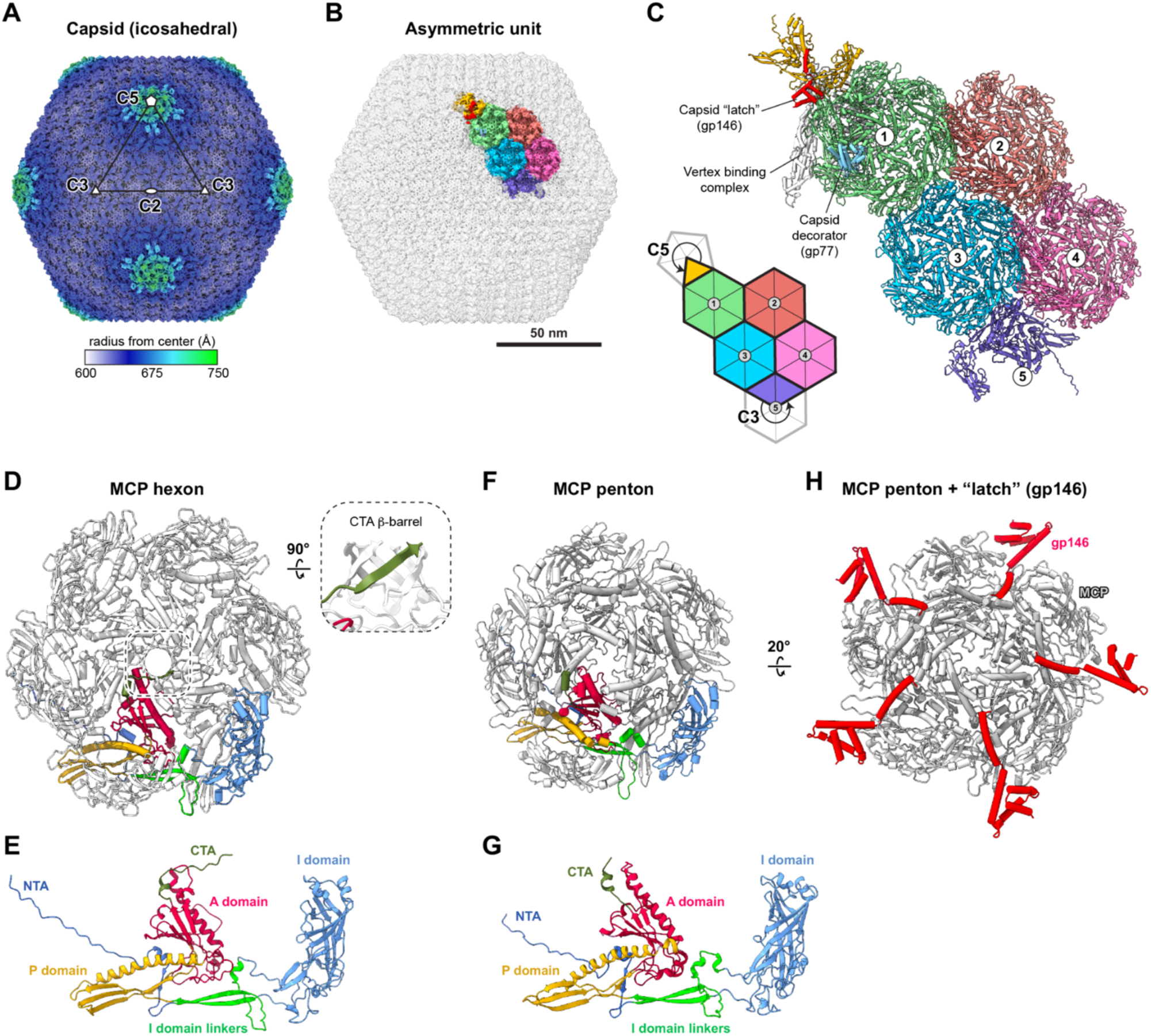
Goslar capsid architecture. A. 6.3 Å resolution icosahedrally-averaged map of the Goslar capsid, colored by radius from center. The two-fold (C2), three-fold (C3), and five-fold (C5) symmetry axes are labeled. B. Icosahedrally-average map of the Goslar capsid with a single asymmetric unit colored. C. Detail view of the Goslar capsid asymmetric unit, with inset diagram. The numbers 1-5 designate MCP hexons (hexon 5 is at a C3 symmetry axis). In addition to MCP, the capsid “latch” protein (gp146) is shown in red; the capsid decorator protein (gp77) is shown in light blue bound to hexon 1, and the vertex binding complex is shown in white. D. Cartoon view of an MCP hexon, with one MCP protomer colored by domain as in panel (e). Inset: rotated view of the six-stranded β-barrel formed by the CTA at the center of the hexon. E. Cartoon view of an MCP protomer in the hexon, with domains colored and labeled. See **Figure S7D-E** for details of hexon-hexon packing. F. Cartoon view of an MCP penton, with one MCP protomer colored by domain. G. Cartoon view of an MCP protomer in the hexon, with domains colored and labeled. Compared to the conformation in the hexon, in the penton the I domain undergoes a ∼30° rotation with respect to the other domains (**Figure S7C**). H. View of the MCP penton with capsid “latch” protein (gp146; red) shown.

The MCPs of Goslar and PhiKZ show strong structural homology, with an overall Cα r.m.s.d. of 4.3 Å, despite their relatively low sequence conservation (25.8%) (**Figure S7A-B**). Both Goslar and PhiKZ MCPs show a canonical HK97 fold^43,44^, with an N-terminal arm (NTA), a peripheral domain (P domain), an axial domain (A domain), an insertion domain (I domain), and a short C-terminal arm (CTA; **Figure 2D-E**). MCP protomers in pentons and hexons possess similar overall structure (**Figure S7C**), with each MCP protomer’s I domain packing against the I domain linkers and P domain of a neighboring protomer, and the A domains of all protomers packing on one another at the center of the assembly (**Figure 2D,F**). The extended NTA also forms a large interface with a neighboring MCP protomer, likely contributing to overall capsid stability. In the hexon, the short CTAs form a six-stranded β-barrel at the very center of the assembly (**Figure 2D** *inset*); this structure is not observed in the penton. Overall, each MCP protomer in both the penton and hexon interacts with four other protomers in the assembly. The most significant difference between the MCPs of the hexon and penton is a ∼30° twist/bend of each protomer’s I domain relative to the other domains (**Figure 2E,G, Figure S7C**); this results in a pyramid-like architecture for the penton, compared to the flat architecture of the hexon. The distinctive structure of the penton is stabilized by the “latch” protein gp146 (**Figure 2H**).

Side-by-side packing of MCP hexons (and pentons) is mediated by two sets of interactions. The major interaction involves the symmetric packing of two protomers’ I domains against one another, with residues Y403, S405, R414, and E421 forming a symmetric hydrogen-bond network (**Figure S7D-E**). On either side of this interface, an extended loop comprising residues 211-229 reaches from each protomer to contact a nearby surface on the I domain of the opposite protomer (**Figure S7E**). These interactions also resemble how MCP hexons pack against one another in different phage capsids^43,44^.

### The vertex binding complex

The recent cryoEM structure of the *Pseudomonas*-infecting chimallivirus PhiKZ capsid revealed a “vertex binding complex” (VBC) comprising seven proteins that form an interdigitated assembly at the inner surface of each non-portal vertex, docked on the inside of the central MCP penton^41^. We similarly observe a five-fold symmetric multiprotein complex docked on the inner surface of the vertex (**Figure 3A-C**), comprising five proteins: gp42, gp59, gp65, gp66, and gp192 (**Figure 3D**). Four of these proteins have clear homologs in the PhiKZ vertex binding complex (**Figure 3E**), including gp42 (PhiKZ gp119), gp59 (PhiKZ gp91), gp65 (PhiKZ gp86), and gp66 (PhiKZ gp85). Goslar gp65 is much longer than its PhiKZ homolog (gp86), and includes a second folded domain not present in PhiKZ gp86 that occupies the same space as PhiKZ gp28 (**Figure 3D-E**). Goslar does encode a predicted homolog of PhiKZ gp028 (Goslar gp219), but we do not observe this protein in our cryoEM map. We also do not observe homologs of two minor PhiKZ vertex-binding complex proteins, gp162 and gp184.

**Figure 3.**
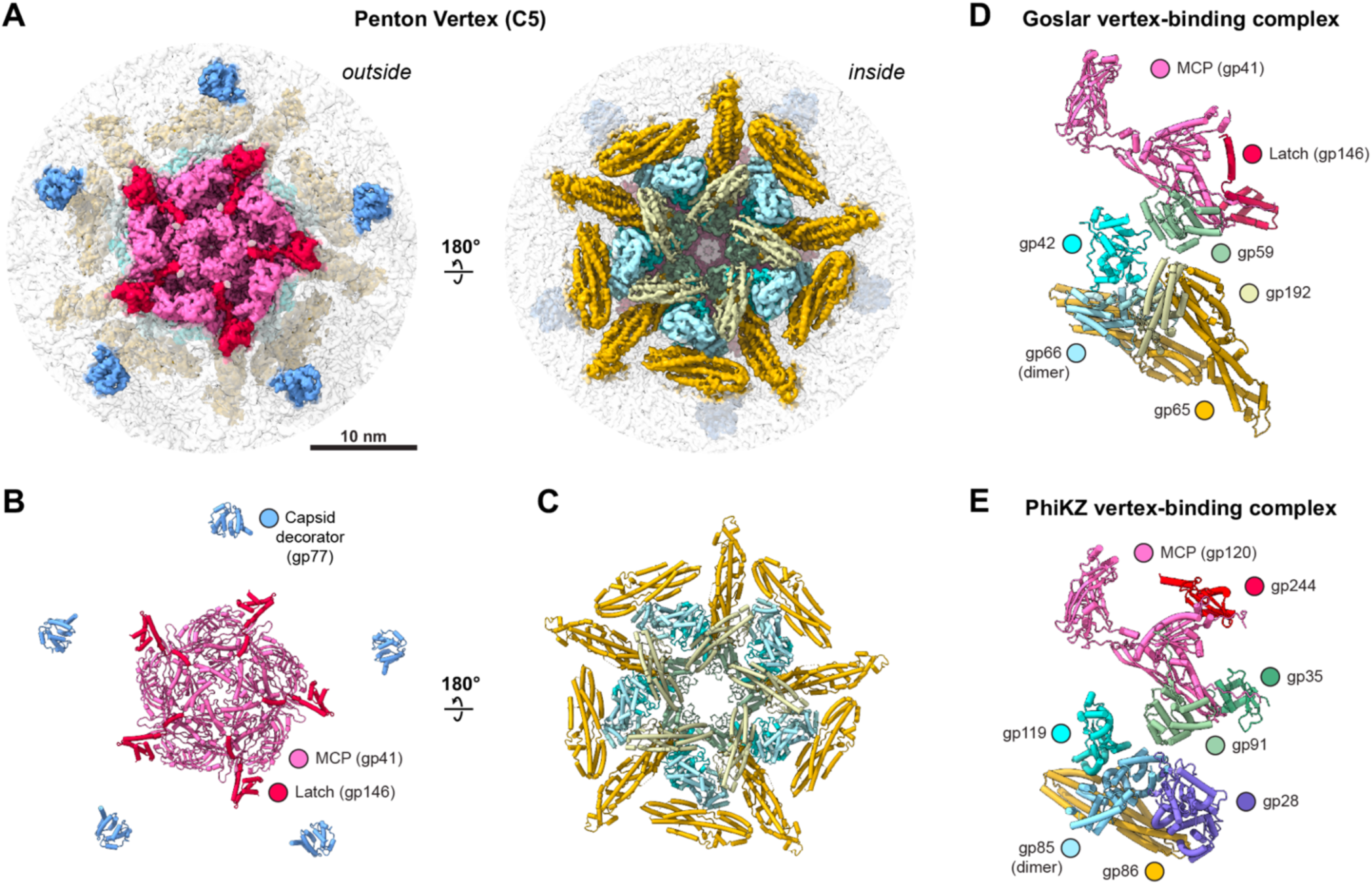
The vertex binding complex. A. Two views of the 3.57 Å resolution C5 penton map. *Left:* view from outside the capsid, with density corresponding to the central MCP penton colored pink (surrounding MCP protomers transparent), the latch protein (gp146) red, and the capsid decorator protein (gp77) blue. *Right:* view from inside the capsid, with vertex binding complex colored as in panel (C). B. Cartoon view of the MCP, latch, and decorator proteins as shown in panel (A). C. Cartoon view of the vertex binding complex as shown in panel (A). D. One unit of the vertex-binding complex, with proteins colored as in panels (A)-(C) and labeled. E. One unit of the vertex-binding complex from the *Pseudomonas* phage PhiKZ (PDB 8Y6V)^41^, with proteins homologous to Goslar vertex-binding complex proteins colored identically to their homologs in panel (D). For clarity, minor components of the PhiKZ vertex-binding complex without Goslar homologs (gp162 and gp184) are not shown. PhiKZ gp35 (dark green) decorates the outer surface of the PhiKZ capsid, and does not possess a Goslar homolog.

In addition to the above confidently-assigned proteins, we observed additional density at low contour surrounding the vertex binding complex and packed against the capsid (**Figure S8**). This density resembles an extended α-helical bundle reminiscent of the PF12699 fold, of which there are several encoded by Goslar (see below). We were unable to confidently assign this density to any Goslar protein.

### Architecture of empty capsids

In addition to intact Goslar virions, we also determined a 5.2 Å resolution icosahedrally-averaged structure of the empty Goslar capsid (**Figure S4A-B**). Compared to the intact virion capsid, the empty capsid has an overall identical symmetry and arrangement of MCP protomers (**Figure S4C-H**). Empty capsids also possess the penton-stabilizing “latch” protein (gp146) and the capsid decorator protein gp77. The most significant differences compared to intact virion capsids are that empty capsids lack both an identifiable portal and also lack the internal vertex binding complex (**Figure S4D,G**). These observations suggest that empty capsids represent assemblies that form without a portal, and are therefore fully symmetric, closed shells unable to be filled with genomic DNA and attached to tails. The lack of the vertex binding complex in empty capsids also suggests that the proteins comprising this complex are not integral structural components of the capsid; rather, they may be packaged after capsid assembly, perhaps concomitant with genome packaging.

### Portal/Neck architecture

Linking capsids to tails is the portal/neck complex, which comprises six different proteins. We determined a 3.8 Å structure of the C12-symmetric portal and upper neck (**Figure 4A** panels i-ii), and a 4.2 Å resolution structure of the C6-symmetric lower neck (**Figure 4A** panels iii-iv). The C12-symmetric region (**Figure 4B**) comprises 12 copies each of the portal protein gp36, the adaptor protein gp233, and the N-terminal domain of the stopper protein gp50. The portal protein is positioned at the top of the assembly, with 12 copies arranged in a ring around a central channel that measures ∼74 Å at the top, decreasing to ∼60 Å at the bottom of the portal. The portal protein itself shares a similar overall architecture with other phage portal proteins, with the large wing domain, stem and clip domains extending downward from the wing domain, and C-terminal crown domain at the top of the assembly (**Figure 4B**). The wing domain spans residues 5-519 and 643-777 of gp36, and the stem and clip domains span residues 520-642. These domains contact the adaptor protein gp233, which in turn packs against the N-terminal domain of the stopper protein gp50. The crown domain of gp36 spans residues 778-923, and reaches counter-clockwise (as viewed from above) across the top of the portal assembly to pack against the crown domains of three neighboring protomers. The portal protein gp36 and the adaptor protein gp233 show structural similarity to the T4 portal (gp20) and neck (gp13) proteins, respectively. Whereas the T4 gp13 dodecamer contacts the hexameric protein gp14, however, the Goslar adaptor protein gp233 contacts the dodecameric stopper protein gp50 (see below).

**Figure 4.**
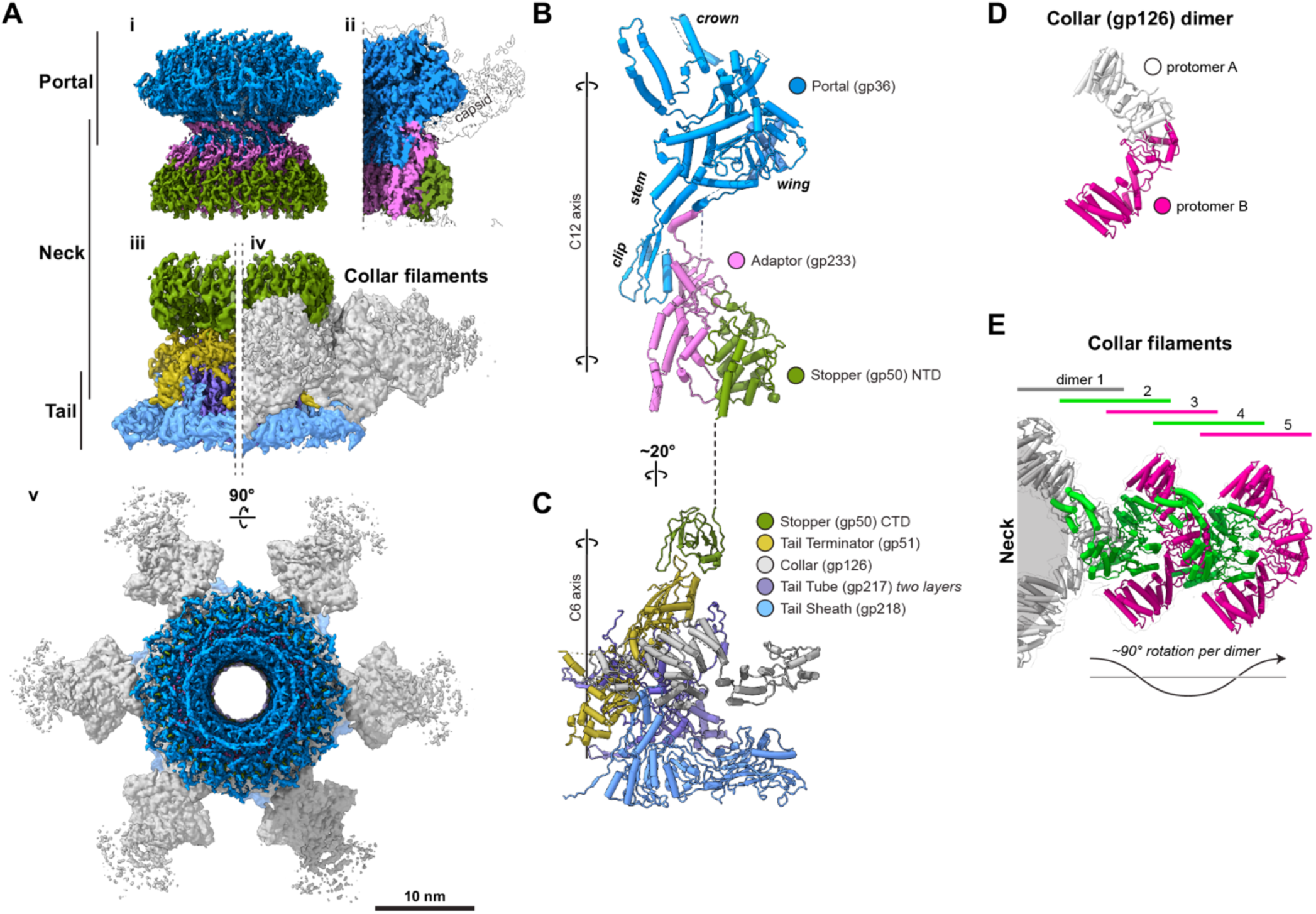
Portal and neck architecture. A. Refined cryoEM maps of the Goslar portal and neck structures. (i) 3.82 Å resolution map of the portal and upper neck (C12 symmetry) colored by protein; (ii) cutaway view of C12-symmetric map with density corresponding to the surrounding MCP capsomers shown in white; (iii) 4.16 Å resolution map of the lower neck (C6 symmetry) colored by protein, with density for the collar subtracted; (iv) Lower neck map showing density for the collar and collar filaments in gray; (v) top-down view of combined portal/neck maps. B. Cartoon view of the C12-symmetric portal and upper neck, with the portal protein (gp36) blue and domains labeled, adaptor protein (gp233) pink, and stopper protein (gp50) green. Only the N-terminal domain of the stopper protein is shown. C. Cartoon view of the C6-symmetric lower neck, with stopper protein (gp50) green, tail terminator (gp51) yellow, collar protein (gp126) gray, tail tube protein (gp217; two layers shown) purple, and tail sheath protein (gp218) blue. Only the C-terminal domain of the stopper protein is shown. Only the inner-most dimer of the collar protein is shown. D. Cartoon view of one dimer of the collar protein (gp126), with one protomer white and the second protomer magenta. E. Top-down view of one collar filament, with the neck at left, and five dimers of the collar protein colored (from inside to outside) gray, green, magenta, green, and magenta.

Below the C12-symmetric upper neck is the C6-symmetric lower neck (**Figure 4C**). The stopper protein (gp50) mediates this symmetry switch, showing C12 symmetry in its upper N-terminal domain that contacts the adaptor protein gp233, and a C6-symmetric alternating inside-outside conformation in its lower C-terminal domain. The C-terminal domain of the stopper protein binds the hexameric tail terminator protein gp51, which in turn binds the top layers of both the inner tail tube protein gp217 and the outer tail sheath protein gp218 (discussed below). Finally, the tail terminator protein also interacts with the “collar” protein gp126. This protein forms a homodimer (**Figure 4D**), and six gp126 homodimers form a ring around the tail terminator protein hexamer (**Figure 4C**). Each gp126 homodimer in this collar structure serves as a docking point for a helical array of gp126 dimers that extends outward from the collar (**Figure 4E**). We could confidently visualize five gp126 homodimers in each of these “collar filaments”, with more visible at low contour levels in our map.

The portal/neck architecture of Goslar closely resembles that of PhiKZ, with the major exception that PhiKZ entirely lacks the collar filaments we observe around the Goslar neck. Instead, PhiKZ possesses a “whisker ring” comprising 6 copies of gp144 and 48 copies of gp34, for which there are not identified homologs in Goslar^34^. The biological roles of the whisker ring in PhiKZ, and of the collar filaments in Goslar, are not known.

### Tail architecture

The Goslar tail is structurally similar to other contractile-tail phages, and is composed of the tail tube protein (gp217) and the tail sheath protein (gp218) forming a C6-symmetric helical array (**Figure 5A-D**). Gp218 is structurally similar to the tail sheath proteins of other phages^45^, with an N-terminal central domain, an outer sheath domain, a C-terminal domain, and a C-terminal extension (**Figure 5E**). To form the extended sheath structure, the outer sheath domain of one protomer interacts with the central domain of a neighboring protomer in the same layer, with the C-terminal domain of a protomer from a lower layer packing against the inside surface of these domains (**Figure S9A-C**). The tail tube protein gp217 is also structurally related to other tail tube proteins^46^, with a small N-terminal domain, conserved central domain, and C-terminal extension (**Figure 5F**). The tail tube protomers form an extensive interlocking network, with the central domain of each protomer packing against the N-terminal domain of a neighboring protomer in the same layer, and each protomer’s C-terminal extension threading into the two layers above it (**Figure S9D-G**). In our structure of the extended (uncontracted) Goslar tail, the tail tube and sheath adopt the same helical symmetry.

**Figure 5.**
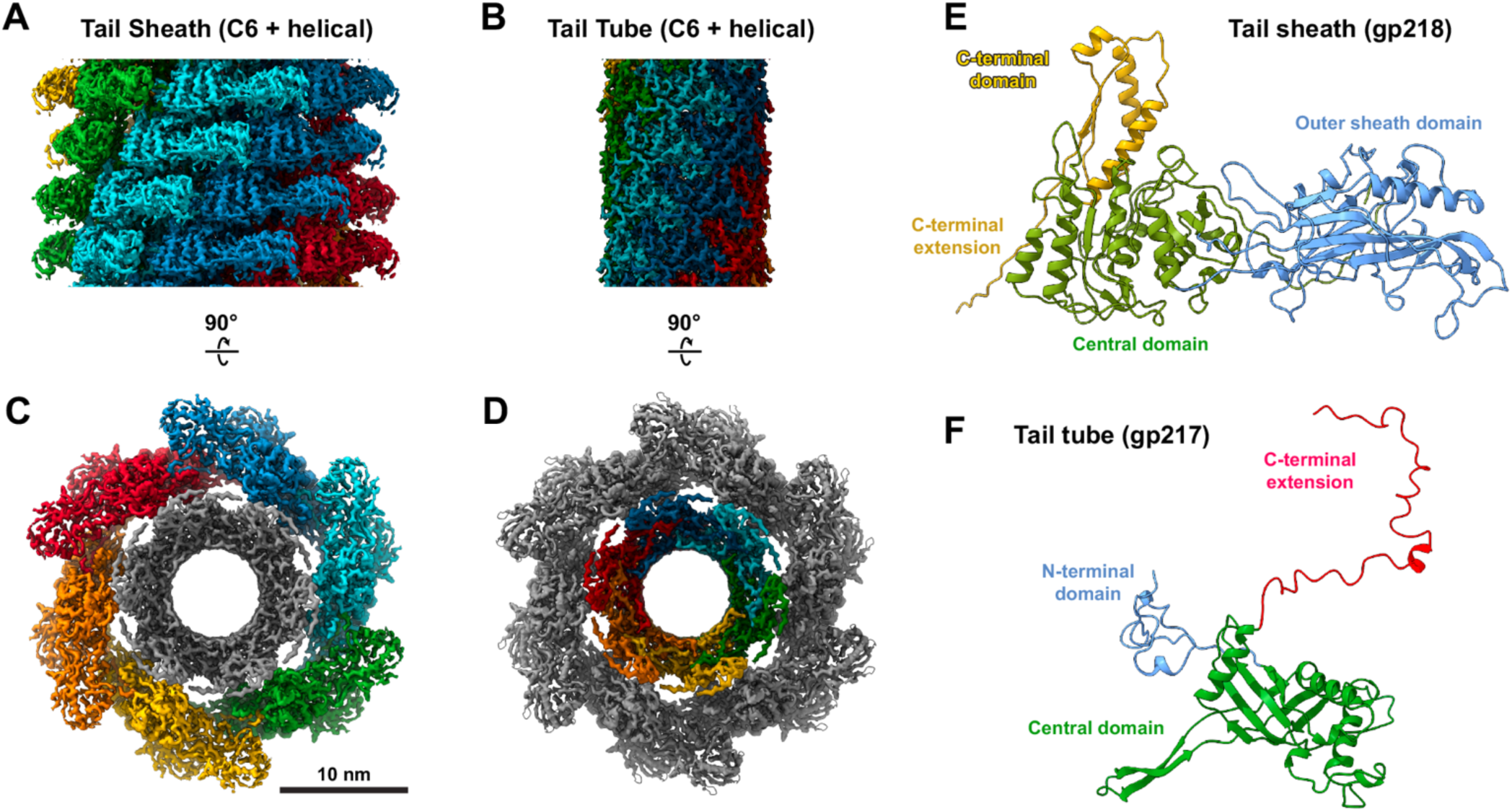
Tail architecture. A. Side view of the tail sheath map, with each helical stack of tail sheath proteins (gp218) colored differently. B. Side view of the tail tube map, with each helical stack of tail tube proteins (gp217) colored differently. C. Top-down view of the combined tail tube/sheath map, with density corresponding to the tail sheath protein (gp218) colored. D. Top-down view of the combined tail tube/sheath map, with density corresponding to the tail tube protein (gp217) colored. E. Cartoon diagram of the tail sheath protein (gp218), colored by domain. F. Cartoon diagram of the tail tube protein (gp217), colored by domain.

In the intact Goslar virion, there are 47 layers of the tail sheath protein, and 46 layers of the tail tube protein, similar to recent findings for PhiKZ (44 layers of tail sheath plus 43 layers of tail tube)^34^. At the neck, the uppermost layer of the tail tube protein is capped by the tail terminator protein (gp51; **Figure 4C**). Gp51 also packs against the upper-most layer of the tail sheath protein, which is adjacent to the second layer of the tail tube protein (**Figure 4C**). At the baseplate (see next section), the lower-most layer of the tail tube protein is capped by the two-protein tail tube initiator complex: gp62 which is homologous to gp217, and gp203 which binds both gp62 and the spike cap protein gp48 (**Figure 6A-B**). The two-layer tube initiator complex is surrounded by the final two layers of the tail sheath protein, which pack against gp172 and gp63 in the inner baseplate (**Figure 6C**).

**Figure 6.**
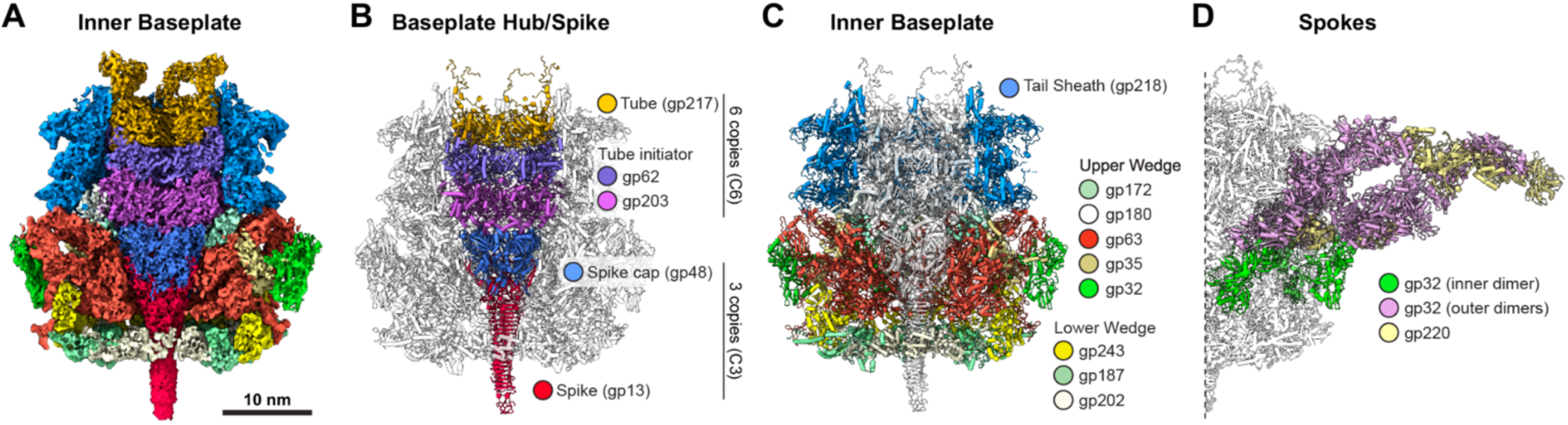
Baseplate architecture. A. 3.88 Å resolution cryoEM map of the Goslar baseplate hub/spike and inner baseplate/wedges, colored by protein as indicated in panels (B) and (C). B. Cartoon view with baseplate hub/spike proteins colored, and inner baseplate/wedge proteins shown in white. C. Cartoon view with the inner baseplate/wedge proteins colored, and hub/spike proteins shown in white. D. Cartoon view of the spokes connecting the inner and peripheral baseplates. Two spokes are shown, each with the inner-most dimer of gp32 (also shown in panel (B)) green, the outer four gp32 dimers pink, and two gp220 protomers light yellow. See **Figure S10A-D** for details on spoke architecture, and **Figure S10E-F** for side and top views of the entire baseplate.

The related chimallivirus PhiKZ has been directly shown to have a contractile tail, with the contracted tail sheath measuring about half the length of the extended tail tail sheath^32,39^. Similarly, we observe a few Goslar particles with contracted tails (**Figure S9H**). These particles were too few in number to perform structure determination on the contracted tails.

### Baseplate architecture

The Goslar baseplate is a large structure measuring nearly 100 nm across (**Figure 6, Figure S10**), and is made up of three main elements: (1) the baseplate hub/spike; (2) the inner baseplate; and (3) the peripheral baseplate. The inner and peripheral baseplates are connected by six “spokes,” creating an overall architecture that resembles a wheel.

From our 3.9 Å resolution map of the baseplate (**Figure 6A**), we were able to confidently build atomic models for the baseplate hub/spike (**Figure 6B**) and inner baseplate/wedges (**Figure 6C**). The baseplate hub comprises the spike protein gp13 (three copies, C3 symmetry), the spike cap protein gp48 (three copies, C3 symmetry), and the tail tube initiator complex (gp203 and gp62, six copies each, C6 symmetry). The bottom layer of the tail tube protein gp217 sits atop gp62 (**Figure 6B**). The architecture of the baseplate hub/spike closely resembles that of phage T4, which also possesses a central spike (gp5 and gp5.4), a spike cap (gp27), and a two-layer tail initiator complex (gp48 and gp54) that links to the tail tube protein (gp19)^47^.

The baseplate hub is surrounded by a six-fold symmetric inner baseplate structure (also termed the “wedges” by analogy with T4) comprising seven different proteins, that we assigned to “upper” and “lower” substructures (**Figure 6C**). The upper wedge comprises four proteins (gp172, gp63, gp35, and gp32) and the lower wedge comprises three proteins (gp243, gp187, and gp202). Our cryoEM map allowed us to confidently assign all density of the baseplate hub/spike and inner baseplate/wedges, and build and refine atomic models for all proteins.

Extending outward from the inner baseplate/wedges are six “spokes” that connect to the peripheral baseplate, which is poorly resolved in our 3.9 Å map. We performed a focused refinement using a mask covering 1/6th of the spoke and peripheral baseplate region, resulting in a 7.9 Å resolution map (**Figure S6D-E**). By careful analysis of AlphaFold-predicted structures and predicted protein-protein interactions, we determine that the spoke comprises a filament of gp32 bound to gp220. Gp32 is predicted to assemble into a two-fold symmetric homodimer, and we could confidently build and refine one gp32 homodimer alongside gp35 in the inner baseplate/wedge (**Figure 6D, Figure S10A-D**).

AlphaFold predicts that gp32 can also form a filament through end-to-end assembly of homodimers, and using the resulting predicted filament structure, we could place four additional gp32 homodimers (ten copies total) extending outward to connect the inner and peripheral baseplates. Finally, we could also tentatively place two copies of gp220 against the gp32 filament, using an AlphaFold-predicted structure of a complex of these two proteins. Thus, the spokes comprise a repeating structure of gp32 and gp220. Our maps did not allow us to confidently assign any other density in the peripheral baseplate.

Our structure of the Goslar baseplate shows strong similarity to the recent structure of the PhiKZ baseplate in the context of the intact PhiKZ virion^34^. The structures of the baseplate hub/spike and inner baseplate/wedges closely match between these related two phages. In agreement with our structure, each of the inner-peripheral baseplate spokes in PhiKZ comprise five dimers of the gp32 homolog gp139, plus two copies of the gp220 homolog gp27. In addition to these proteins, the higher resolution of the PhiKZ baseplate structure allowed modeling of several additional proteins, all of which have clear homologs in Goslar (**Table S3**). Even so, major portions of the peripheral baseplate in both PhiKZ and Goslar remain unmodeled, leaving the detailed architecture of the baseplate and any conformational changes associated with host recognition and genome injection as open questions for future work.

### Mass spectrometry of Goslar virions identifies abundant virion-associated proteins

Chimalliviruses’ distinctive lifestyle, notably their mechanism of infection that involves injection of their genomic DNA into a lipid-enclosed vesicle, demands that these phages package and inject a suite of proteins alongside their DNA to mediate a successful infection. Chimalliviruses are known to inject the multisubunit virion RNA polymerase (vRNAP)^17–19^; high-copy proteins thought to make up the inner body (e.g. PhiKZ gp93 and its Goslar homolog gp58)^12,13,48^; and other proteins with unknown functions^13^. To identify the full set of proteins associated with the Goslar virion, we subjected our purified Goslar virions to liquid chromatography-tandem mass spectrometry (LC-MS/MS) with label-free quantitation (LFQ). Calibrating the MS2 peak area assigned to peptides from each protein with the known copy number of the MCP and tail tube/sheath proteins we could roughly estimate (+/− about 50%) the per-virion copy number of all proteins detected in our dataset (see **Methods**) (**Figure 7A-B, Table S4**). By this measure, we detected 12 proteins present in >200 copies per virion that were not part of our atomic model of the Goslar virion (**Figure 7B**). Two of these proteins were gp53 (∼330 copies per virion) and gp58 (∼2150 copies per virion), both of which contain a predicted PF12699 domain and are related to several PhiKZ proteins with the same domain (see below). The other ten proteins we detect in >200 copies per virion (from highest to lowest abundance: gp23, gp158, gp214, gp88, gp22, gp208, gp61, gp209, gp7, and gp168) all lack a clearly predicted function from sequence or comparisons of their AlphaFold-predicted structures (**Figure S11**) to known proteins.

**Figure 7.**
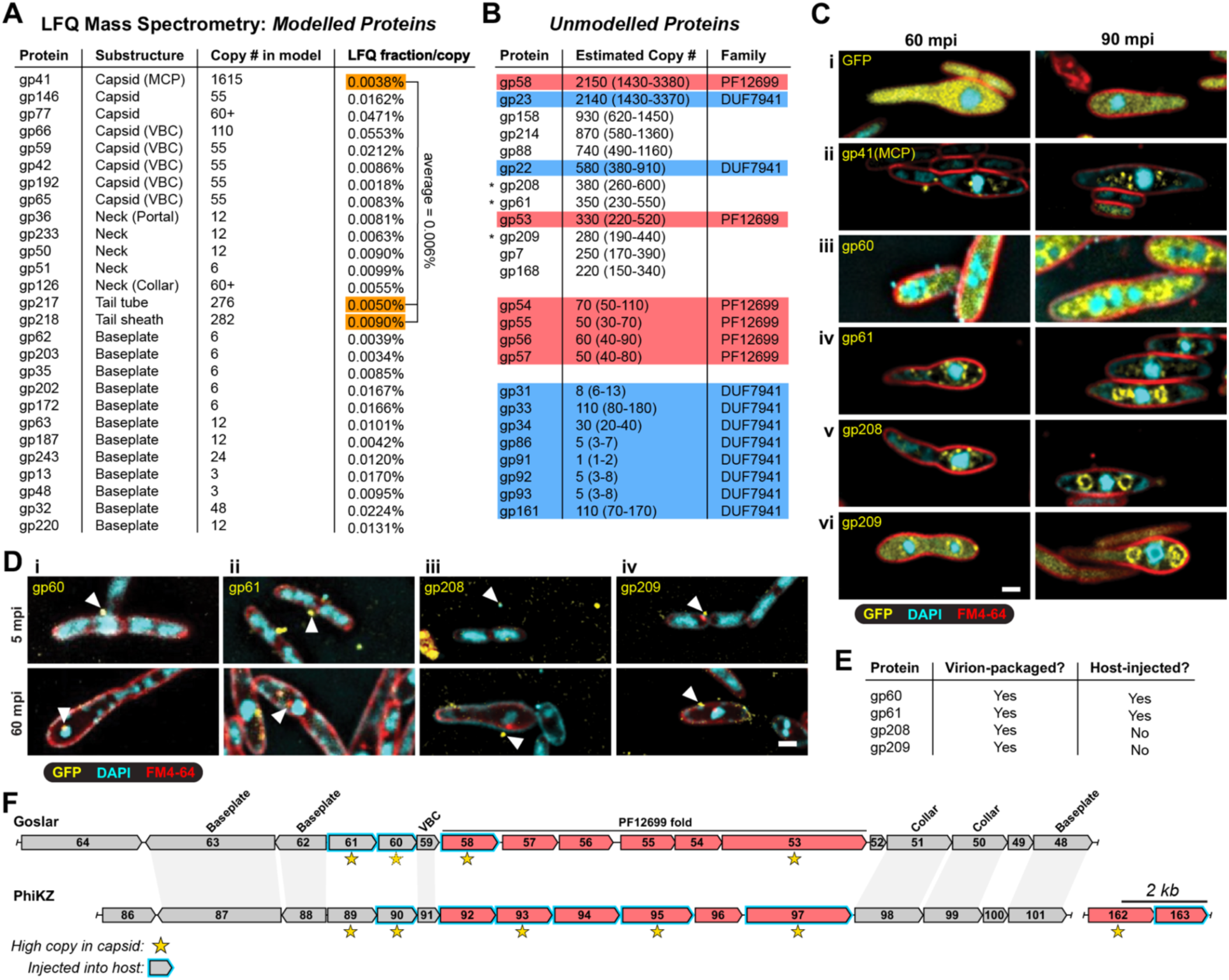
Mass spectrometry identifies abundant virion-associated proteins. A. Results from label-free quantification (LFQ) mass spectrometry analysis of purified Goslar virions, showing 27 structural proteins (gp180 was not detected by mass spectrometry), their assigned substructure, the number of copies modelled, and the LFQ fraction per copy calculated from mass spectrometry. Three proteins with known high copy numbers (highlighted orange: gp41/MCP, gp217/tail tube, and gp218/tail sheath) were used to estimate copy number per virion of unmodelled proteins. See **Table S4** for details. B. Select unmodelled proteins detected by mass spectrometry, and their estimated copy number per virion. Shown is the estimated copy number as calculated using an average LFQ fraction/copy of 0.006% (average of gp41, gp217, and gp218 LFQ/copy), and the range defined by a low estimate using 0.009% (gp218 LFQ/copy) and a high estimate using 0.0038% (gp41 LFQ/copy). The top 12 most-abundant proteins are shown, which includes two PF12699 proteins (highlighted red: gp58 and gp53) and two DUF7941 proteins (highlighted blue: gp23 and gp22). Estimated copy number for all other PF12699 and DUF7941 proteins are also shown. See **Table S4** for details. C. Subcellular localization of GFP and indicated GFP-tagged Goslar proteins in Goslar-infected *E. coli* MC1000 cells, 60 and 90 minutes post infection (mpi). (i) Localization of GFP; (ii) Localization of gp41-GFPmut1 (major capsid protein; MCP); (iii) Localization of gp60-GFPmut1; (iv) Localization of GFPmut1-gp61; (v) Localization of GFPmut1-gp208; (vi) Localization of gp209-GFPmut1; the 60 mpi panel shows a double-infected cell with two phage nuclei. GFP is yellow; DAPI (DNA) is cyan; and FM4-64 (membranes) is red. Scale bar = 1 µm. See **Figure S12** for separate fluorescent channels and DIC images. D. Fluorescence microscopy of Goslar virions with GFP-tagged proteins, 5 and 60 minutes post infection (mpi): (i) gp60-GFPmut1; (ii) GFPmut1-gp61; (iii) GFPmut1-gp208; (iv) gp209-GFPmut1. Arrowheads indicate the location of virions (on the cell surface) or EPI vesicles (within cells). GFP is yellow; DAPI (DNA) is cyan; and FM4-64 (membranes) is red. Scale bar = 1 µm. See **Figure S13** for separate fluorescent channels and DIC images. E. Summary of findings for virion incorporation and host injection for gp60, gp61, gp208, and gp209. F. Schematic of the syntenic genomic regions of Goslar (top; NCBI RefSeq NC_0481701.1) and PhiKZ (bottom; NCBI RefSeq NC_004629.1) encoding a cluster of structural proteins and PF12699 proteins (colored red). Known-homologous genes^25^ are indicated by gray shading, and the substructure to which each Goslar protein is assigned (if any) is noted above each gene. Yellow stars indicate proteins present at high copy number in virions, as determined by mass spectrometry of Goslar virions (this study) and PhiKZ virions^40^. Proteins from either Goslar or PhiKZ that have been demonstrated to be injected into the host are indicated with blue outlines. See **Figure S14** for AlphaFold 3 predicted structures of all Goslar PF12699 proteins.

To determine whether the high-copy proteins we detect by mass spectrometry are virion-associated, we expressed GFP fusions of three high-copy proteins (gp61, gp208, and gp209) plus a fourth protein (gp60; present at an estimated ∼140 copies per virion) that is predicted by AlphaFold to bind gp61 (**Figure S11I**), and examined their localization in Goslar-infected cells. The localization of all four proteins resembled that of the major capsid protein gp41, with puncta surrounding the phage nucleus at 60 minutes post infection (mpi), and puncta clustering at the phage “bouquets” at 90 mpi (**Figure 7C, Figure S12**). We next generated phage lysates from cells expressing each GFP fusion and infected wild-type MC1000 cells to test whether the proteins are injected into host cells. We found that all four proteins showed clear puncta representing Goslar virions (**Figure 7D**). We further found that gp60 and gp61, but not gp208 or gp209, were injected into host cells, co-localized with EPI vesicles in early infections (15 mpi), and formed a single focus on the phage nucleus in late infections (60 mpi) (**Figure 7D, Figure S13**), consistent with prior observations of host-injected proteins in Goslar and PhiKZ^12,13,42,48^. Overall, these data show that gp60, gp61, gp208, and gp209 are all virion-packaged, and that gp60 and gp61 are injected into the host cell along with the phage genome (**Figure 7E**).

In PhiKZ, a set of proteins with a common predicted fold characterized by the Pfam PF12699 domain are present at high copy number in the capsid, and have been proposed to form a large internal assembly termed the “inner body”^38–42^. These proteins (PhiKZ gp92, gp93, gp94, gp95, gp96, gp97, gp162, and gp163) are mostly encoded in a single operon (**Figure 7F**), and most are also injected into the host cell upon infection^13,42,48^. The Goslar proteins gp53 and gp58 also possess a predicted PF12699 domain, and are encoded in a region of the Goslar genome that is syntenic with the PhiKZ region that encodes gp92-gp97 (**Figure 7F**). Like several PhiKZ PF12699 proteins, we also previously showee that Goslar gp58 is injected into the host cell^12^. We performed AlphaFold structure predictions of the proteins encoded near gp53 and gp58 in the Goslar genome, and found that gp54, gp55, gp56, and gp57 share the predicted PF12699 fold (**Figure S14**). We detect all of these proteins in our mass spectrometry: gp53 at ∼330 copies per virion, gp54 at ∼70 copies, gp55 at ∼50 copies, gp56 at ∼60 copies, gp57 at ∼50 copies, and gp58 at ∼2150 copies (**Figure 7B, Table S4**). Thus, while PhiKZ has several similar proteins at ∼100-200 copies per virion^40^, Goslar has a single dominant PF12699 family member in gp58. The biological role of the injected PF12699 proteins, including whether their primary functions are within the capsid or in the EPI vesicle after injection into a host cell, is not known.

We noticed that two of the highest-abundance proteins in the Goslar virion, gp22 (∼580 copies per virion) and gp23 (∼2140 copies), are paralogs of one another and share a common DUF7941 domain (DUF: domain of unknown function) (**Figure S11A,E**). The annotated DUF7941 domain spans the entire length of gp22, which is predicted to comprise an N-terminal globular domain, a disordered linker, and a C-terminal globular domain (**Figure S11E**). Gp23 shares the N-terminal globular domain and disordered linker, but lacks gp22’s C-terminal globular domain. We could not identify any confident structural matches to gp22 or gp23 using the DALI^49^ or FoldSeek^50^ servers. Using HMM searches with the SeqHub web service^51^ and AlphaFold structure predictions, we identified eight other Goslar proteins containing the DUF7941 module: gp31, gp33, gp34, gp86, gp91, gp92, gp93, and gp161 (**Figure 8A**). In three of these (gp86, gp92, and gp161), an N-terminal DUF7941 module is followed by predicted carbohydrate binding, glycoside hydrolase, and/or leucine-rich repeat domains, suggesting a role for these proteins in host recognition (**Figure 8A, S15**). The DUF7941 module is found exclusively in chimalliviruses; for example, PhiKZ is predicted to possess at least three DUF7941 proteins (gp126, gp127, and gp143), and the *Erwinia* phage RAY is predicted to possess at least five (gp8, gp9, gp11, gp215, and gp218).

**Figure 8.**
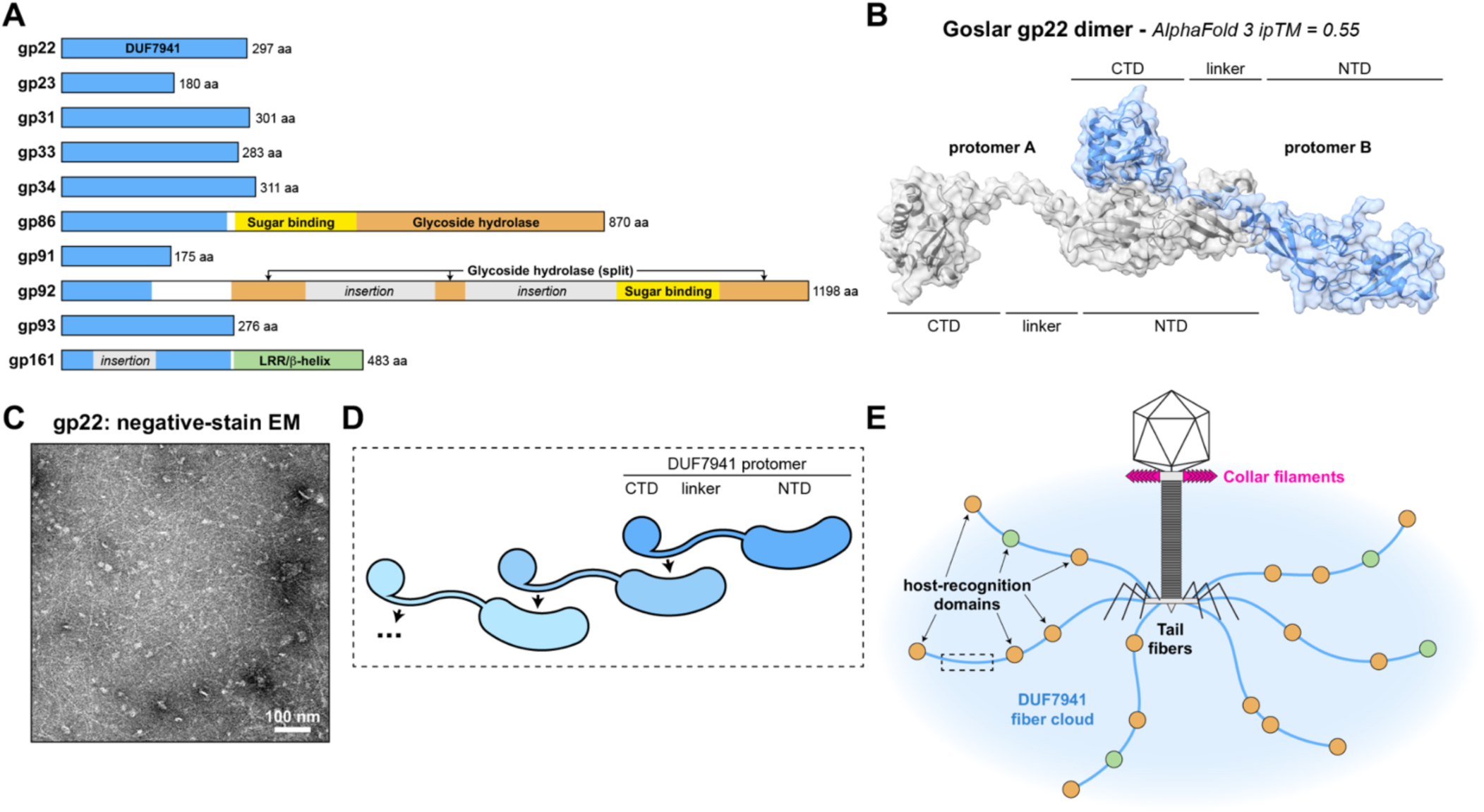
Identification of DUF7941 proteins in Goslar and model for oligomerization. A. Domain schematics of 10 identified DUF7941 proteins in Goslar. DUF7941 domains are shown in blue, carbohydrate-binding domains in yellow, glycoside hydrolase domains in orange, leucine-rich repeat (LRR)/β-helix domains in green, and unknown domains in gray. AlphaFold predictions of all proteins are shown in **Figure S15**. B. AlphaFold 3 predicted structure of a Goslar gp22 dimer. The disordered linker and CTD of one protomer (blue) is predicted to dock against the NTD of a second protomer (gray) in a head-to-tail orientation. See **Figure S16** for further details, and for predicted structures of a Goslar gp23 dimer. See **Figure S17** for the predicted structure of a PhiKZ gp126-gp127 heterodimer, and fitting of that model into cryoEM density of the PhiKZ peripheral baseplate. C. Negative-stain electron microscopy of purified Goslar gp22, showing extended filaments. See **Figure S18** for additional images. D. Model for head-to-tail assembly of DUF7941 domains. E. Speculative model showing likely host-recognition elements in Goslar, with collar filaments shown in magenta, tail fibers in black, and the DUF7941 fiber cloud in blue, with associated carbohydrate binding/hydrolysis domains in orange and LRR/β-helix domains in green.

Curiously, AlphaFold predictions suggest that Goslar gp22 could form head-to-tail oligomers through binding of one protomer’s disordered region to the N-terminal globular domain of another protomer (**Figure 8B, S16A-B**). While Goslar gp23 lacks the C-terminal globular domain of gp22, it is also predicted to form head-to-tail oligomers (**Figure S16C-E**). We used AlphaFold to predict homo- or hetero-oligomer formation between each pair of DUF7941 proteins in Goslar, and identified a network of potential interactions, suggesting that these proteins may form extended, heterogeneous filaments (**Figure S16F**). Relatedly, we found that the PhiKZ DUF7941 proteins gp126 and gp127 are confidently predicted by AlphaFold to form a head-to-tail dimer (**Figure S17A-C**). In a recent structure of the intact PhiKZ virion, the DUF4391 protein gp127 is partially modeled adjacent to the outermost dimer of gp139 (homologous to Goslar gp32)^34^. In the published PhiKZ structure, only the gp127 NTD is modeled (**Figure S17D**), but we could model a full gp126-gp127 heterodimer into unexplained density in the PhiKZ outer peripheral baseplate map (**Figure S17E-F**). Therefore, while no density for DUF4391 proteins is observable in our less well-defined maps of the Goslar peripheral baseplate (**Figure S10**), we propose that chimallivirus DUF7941 proteins universally associate with the peripheral baseplate.

To test whether DUF7941 proteins can form oligomers, we expressed and purified Goslar gp22, and examined the purified protein by negative-stain electron microscopy. We observed filaments and filament bundles up to several hundred nanometers in length (**Figure 8C, Figure S18**). Combined with our AlphaFold predictions suggesting that DUF7941 proteins form head-to-tail oligomers, we propose that chimallivirus DUF7941 proteins form a “cloud” of extended, flexible filaments that facilitate initial host recognition (**Figure 8D-E**). Based on the overall copy number estimated from mass spectrometry (over ∼2500 DUF7941 domain proteins per Goslar virion), protomer spacing predicted by AlphaFold (∼7 nm per protomer), and the likely number of peripheral baseplate anchoring sites (6), we calculate an average DUF7941 filament length of nearly 3 microns in Goslar. Thus, the DUF7941 filament cloud would be expected to dramatically increase the ability of each virion to encounter and bind host cells.

In all, we identified 87 Goslar proteins in our mass spectrometry dataset with a predicted copy number of five or more per virion. These proteins include all five vRNAP subunits (gp78, gp196, gp205, gp207 and gp210), and several proteins annotated as tail or baseplate proteins that likely represent components of the peripheral baseplate/tail fibers (e.g. gp25, gp29, and gp90; **Table S4**). We also detected the nuclear shell protein chimallin (gp246) and proteins known to be chimallin-associated (e.g. gp176/ChmC), plus the tubulin-like protein PhuZ (gp234). These data show that our phage isolation method (including PEG precipitation and CsCl gradient centrifugation) also captured large protein assemblies including phage nucleus fragments and PhuZ filaments.

## DISCUSSION

Here we use cryoEM to define the overall molecular architecture of the chimallivirus Goslar, with our final model comprising 2,888 polypeptide chains from 28 different structural proteins. Our maps of individual phage substructures ranged from ∼3.2 to ∼4.2 Å in resolution, allowing us to confidently build and refine atomic models for most structural proteins. The assistance of modern artificial-intelligence tools including AlphaFold^37^ and ModelAngelo^36^ was crucial for initial assignment of low-resolution density, initial model building, and predictions of protein-protein interactions and complex architectures within each substructure. Overall, the architecture of Goslar is consistent with that of the related phage PhiKZ^34^ and with other tailed phages, with a T=27 icosahedral capsid, 12-fold symmetric portal, helical tail assembly, and baseplate with central spike (three-fold symmetric), inner baseplate/wedges (six-fold symmetric), and an extended peripheral baseplate (six-fold symmetric). While we could assign density for the spokes connecting the inner and peripheral baseplates, our maps were unable to resolve the full peripheral baseplate and tail fibers.

Our cryoEM model and quantitative mass spectrometry analysis of purified Goslar virions reveal several unique properties of this large-genome phage. First, we define a novel assembly of “collar filaments” extending from the portal-neck region, made up of an extended array of gp126 dimers. The role of these filaments is not known: they could act as an environmental sensor akin to the phage T4 whiskers^52,53^, or interact with host flagella similarly to the capsid fibers of phage 7-1-1 (ref. ^54^). There is strong evidence supporting the idea that PhiKZ and related chimalliviruses use the host flagellum for initial recognition, including the observation that these phages tend to infect near the cell pole^55^ and that phage-resistant mutants often affect the flagellum^56,57^. Notably, the collar filaments we observe associated with the Goslar neck region are completely distinct from the “whisker ring” recently reported as surrounding the neck region of the related phage PhiKZ^34^.

Another distinctive feature of Goslar — shared with PhiKZ — is the presence of high-copy virion associated proteins that possess the uncharacterized PF12699 domain. In PhiKZ, these proteins have been proposed to make up the inner body, a large structure that has been observed in the PhiKZ head by low-resolution electron microscopy studies^40–42^, but notably not observed in recent high-resolution studies of the same phage^34^. We do not observe the inner body in our cryoEM micrographs or 3D reconstructions of Goslar, but purified Goslar phages do contain thousands of copies of PF12699 proteins per virion. Gp058 is most abundant, with an estimated 2000+ copies per virion, and we also detect dozens to hundreds of copies of each of the structurally related proteins gp53, gp54, gp55, gp56, and gp57. Many PF12699 proteins (including Goslar gp58) are known to be injected into the EPI vesicle formed upon initial infection of a host^12,13,42,48^. The biological roles of these proteins remain a key unsolved mystery of the chimallivirus life cycle.

Our mass spectrometry also identified a new family of chimallivirus-specific proteins characterized by the DUF7941 domain. We find that Goslar virions incorporate hundreds of copies of the DUF7941 proteins gp22 and gp23, and we also detect significant numbers of eight additional DUF7941 proteins associated with Goslar virions. Our AlphaFold predictions and negative-stain electron microscopy of gp22 indicate that the DUF7941 module can form head-to-tail oligomers. Combined with our finding that several Goslar DUF7941 proteins encode additional domains associated with host recognition, we propose that these proteins associate with the peripheral baseplate and form a “DUF7941 fiber cloud” extending up to several microns from the phage itself.

Our work sets the stage to answer a range of pressing questions regarding chimallivirus infection and development. Aside from determining the relative contributions of different elements — collar fibers, tail fibers, and newly-identified DUF7941 fibers — to host recognition, a key question is how these phages inject their genomes into a lipid vesicle in their host. The tail architecture of Goslar is similar to other tailed phages, and we observe occasional virions with contracted tail sheaths in our cryoEM micrographs (**Figure S9H**). Key next steps will be to determine in detail the structural changes that occur upon tail sheath contraction; how host cells are penetrated, and how the lipid bilayer comprising the EPI vesicle is shaped upon injection of the genome and associated proteins.

## MATERIALS AND METHODS

### Phage Goslar preparation

Bacteriophage Goslar was originally acquired from Johannes Wittman at the DSMZ^28^. For phage propagation, 100 µL of medium-titer viral lysate (10^8^-10^9^ PFU/ml) was incubated with 0.5 mL *E. coli* MC1000 cells for 10 minutes and plated on LB agar plates after mixing with 4.5 mL 0.35% LB top agar, and incubated overnight at 37°C. Lysis of host cells were observed by plate clearing as compared to a control plate containing only MC1000 cells. Viral particles were harvested by addition of 5 mL LB media, incubation overnight at 4°C, followed by gentle scraping using a sterile cell spreader. After collection, lysates were subjected to a low-speed spin (3220 x g for 10 minutes) followed by filtration through a 0.45 µm syringe filter. The resulting supernatant was treated with DNase I (1 µg/mL) and RNase A (1 µg/mL) for 1 hour at 37°C, followed by addition of 2 mL (0.4 mL per 1 mL phage lysate) phage precipitating solution (30% PEG 8000, 3.3 M NaCl) and kept at 4°C overnight. This was followed by centrifugation at 10,000 x g for 20 minutes to sediment phage particles. The pellet was resuspended in phage buffer (10 mM Tris-HCl pH 7.5, 10 mM MgSO_4_, 68 mM NaCl, 1mM CaCl_2_), mixed with solid CsCl (0.55 grams per 1 mL resuspended phages), and spun in an ultracentrifuge (55,000 RPM, 12 hours, using a Beckman-Coulter TLS-55 Rotor). The opalescent phage “band” was extracted with a syringe, buffer-exchanged to phage buffer, and concentrated by ultrafiltration (Amicon Ultra, EMD Millipore) to a final concentration of 10^10^ PFU/ml for mass spectrometry.

### Cryo-EM sample preparation and data collection

Due to low particle density on grids when using CsCl-purified phages, we skipped PEG precipitation and ultracentrifugation for cryoEM samples. Instead, 3 µL of filtered phage lysate was directly applied to Carbon-coated Lacey-Carbon grids (Electron Microscopy Sciences) and imaged using a Titan Krios microscope (ThermoFisher Scientific, operated at 300 kV) equipped with a K3 direct electron detector (Gatan) and a Biocontinuum Energy Filter (Gatan). Three cryoEM datasets were collected. Dataset 1 consisted of 4,224 movies collected in super-resolution mode under a nominal magnification of 33,000x (corresponding to a pixel size of 1.3 Å) with a total dose of 35 e^−^/Å^2^ and a defocus range of −0.5 to −4 µm. Datasets 2 (5,677 movies) and 3 (7,841 movies) were collected in counting mode under a nominal magnification of 64,000x (corresponding to a pixel size of 1.4 Å) with a total dose of 50e^−^/Å^2^ with defocus range of −1 to −3 µm. All data-collection statistics are reported in **Table S1** (Dataset 1) and **Table S2** (Datasets 2 and 3).

### Cryo-EM data processing

#### Intact Goslar virions

All cryoEM processing was performed with cryoSPARC version 4.7 (ref. ^35^). Intact virions were manually picked from Dataset 1, with picking performed such that particles were centered on capsid’s portal vertex. Particles were aligned using 2D classification, then re-centered to the midpoint of the virion along the tail for extraction. Particles were extracted with a 3600 pixel box, then Fourier-cropped to 800 pixels. For 3D reconstruction, 3,033 particles were subjected to Homogenous Reconstruction followed by Local CTF Correction, then Local Refinement, yielding a 19.46 Å resolution map of the intact Goslar virion (**Figure S1**).

#### Empty Capsid

From Dataset 1, ∼100 empty capsids were manually picked and subjected to 2D classification to generate templates for automated particle picking. Template-based picking followed by multiple rounds of 2D classification yielded a final set of 4,535 particles, which were extracted with an 1800 pixel box size and Fourier-cropped to 960 pixels (the maximum particle size possible given the memory limits of the NVIDIA RTX A5000 GPUs used for refinement). After Ab Initio reconstruction, multiple rounds of Homogeneous Refinement and Global CTF correction were performed with icosahedral (I1) symmetry to account for the large particle size. For Homogeneous Refinement, the following non-default parameters were used: Fit Spherical Aberration, Fit Tetrafoil, Fit Anisotropic Mag., and Do EWS correction (**Figure S4**).

To identify the portal vertex, particles were symmetry-expanded, rotated to align particles on the C5 axis, re-extracted with a 600-pixel box, then duplicate particles were removed and a homogeneous reconstruction was performed with a final set of 54,368 symmetry-expanded particles. We then performed 3C Classification with 2-10 3D classes. No portal vertices were identified in empty capsids by this analysis (**Figure S4B**).

#### Capsid

From Datasets 2 and 3, capsids attached to tails were initially picked using the “blob picker” job in CryoSPARC with particle diameter 1000-1800 Å, followed by iterative 2D classifications to define a 5,214 particle set. Particles were extracted with an 1700-pixel box size, Fourier-cropped to 800 pixels, then subjected to global CTF refinement followed by homogenous refinement with icosahedral (I1) symmetry imposed. This yielded a 6.28 Å resolution icosahedral map for the phage capsid (**Figure S2, S3A-B**). Particles were symmetry expanded (I1) and re-centered on either the C3 (hexon) or C5 (penton) symmetry axes using the volume alignment Job in cryoSPARC. For the reconstruction of the C3 hexon map, C3-centered particles were re-extracted with a 480 pixel box size, and duplicate particles were removed. The remaining particles were subjected to local refinement with C3 symmetry, yielding a 3.57 Å resolution map which was iteratively improved by several rounds of local CTF refinement and local refinement (**Figure S2, S3C-D**).

C5-centered particles were used to reconstruct an initial volume with the “homogenous reconstruct only” job in cryoSPARC. The resulting volume was used for a 3D classification job with 14 3D classes, to isolate the unique portal vertex class (see below). Particles assigned to non-portal vertex classes were combined and subjected to iterative local refinement and local CTF refinement with C5 symmetry (**Figure S2, S3E-F**).

#### Portal & Neck

Symmetry-expanded capsid particles assigned to the portal vertex class in 3D classification (see above) were re-extracted with a box size of 640 pixels, then symmetry-expanded (C12) and subjected to local refinement - this represents the upper region of the portal-neck assembly (**Figure S2, S3G-H**). The same particles were symmetry-expanded (C6) and subjected to local refinement - this represents the lower neck region (**Figure S2, S3I-J**).

#### Tail

From Datasets 2 and 3, tail particles were picked using the filament tracer job in cryoSPARC with filament diameter set to 300 Å and particle separation distance set to 150 Å. Initial particle picks were curated by 2D classification, resulting in a final set of 47,640 particles. These particles were extracted with a box size of 420 pixels, followed by helical refinement with helical twist 21.48°, helical rise 39.16 Å, and symmetry order 4 to yield a 3.39 Å tail map (**Figure S5A-C**). Particles were symmetry-expanded, then local refinement was performed with focused masks around either the tail tube or the tail sheath, yielding a tail tube map at 3.22 Å resolution (**Figure S5D-F**) and a tail sheath map at 3.36 Å resolution (**Figure S5G-I**).

#### Baseplate

From Datasets 2 and 3, ∼100 baseplate particles were manually picked and subjected to 2D classification to generate templates for automated particle picking. Automatically-picked particles were curated by 2D classification, yielding a final set of 5,539 baseplate particles. Particles were extracted with a box size of 800 pixels and Fourier-cropped to a box size of 600 pixels, then homogeneous reconstruction was performed with C6 symmetry to obtain an initial 4.35 Å-resolution baseplate map. Density representing the lower tail was subtracted, then particles were symmetry-expanded (C6) and subjected to iterative local refinement and local CTF refinement to yield the final 3.88 Å resolution baseplate map (**Figure S6A-C**). To visualize the spokes and peripheral baseplate, a mask encompassing 1/6th of this region was generated and used as a focus mask with symmetry-expanded particles. This resulted in a 7.88 Å resolution peripheral baseplate map (**Figure S6D-E**).

### Model Building and refinement

Initial atomic models for several proteins were built by ModelAngelo^36^.For well-annotated structural proteins such as the portal, sequence-based ModelAngelo was run to generate an initial model which was validated by Alphafold3 predictions^37^, followed by manual building in COOT^58^ and refinement using PHENIX^59^. For substructures where candidate proteins were unknown, sequence-free ModelAngelo was performed, then the Modelangelo-assigned sequences were compared against the Goslar proteome using BLAST to identify candidate proteins. Candidates were validated by docking AlphaFold3-generated models^37^, and by leveraging predicted protein-protein interactions with neighboring proteins, generated by AlphaFold3 co-predictions. All models were manually rebuilt in COOT^58^ and refined in phenix.refine^59^ with strict non-crystallographic symmetry constraints for symmetry-related protomers; secondary structure restraints; and reference-model restraints (using confident AlphaFold3 models as reference models when available). For the baseplate spoke/peripheral baseplate region, manually-docked AlphaFold models for gp32 (dimers) and gp220 (separate N- and C-terminal domains) were rigid-body refined using the 7.88 Å resolution spoke/peripheral baseplate map. All refinement statistics are reported in **Tables S1-S2**.

A final composite model of the full Goslar virion (**Figure 1A**) was generated by progressively building overlapping large substructures (full capsid with portal, portal with neck, neck with full tail, tail with baseplate) and manually docking these substructures to one another and then to the asymmetric Goslar virion map. Structures were visualized in UCSF ChimeraX^60^.

### Bioinformatics and structure prediction

To define Goslar proteins with PF12699 or DUF7941 domains, we used annotations available at InterPro (https://www.ebi.ac.uk/interpro/)^61^ and examined sequences using SeqHub^51^ (https://seqhub.org; Tatta Bio). For structure prediction of unmodelled proteins including PF12699 and DUF7941 proteins, predictions were performed using AlphaFold3 (ref. ^37^) through the AlphaFold server (https://alphafoldserver.com). Structures were visualized in UCSF ChimeraX^60^, and PAE plots were generated using PAE Viewer (https://pae-viewer.uni-goettingen.de)^62^.

### Mass spectrometry sample preparation

For mass spectrometry, CsCl-purified Goslar phage samples were desiccated and reconstituted in 200 µl of 6M Guanidine-HCl. The samples were then boiled for 10 minutes followed by 5 minutes cooling at room temperature. The boiling and cooling cycle was repeated a total of 3 cycles. Proteins were precipitated with addition of 1.8 mL methanol final concentration, 90% methanol v/v) followed by vortexing and centrifugation at 14000 RPM on a benchtop microcentrifuge for 10 minutes. The soluble fraction was removed by flipping the tube onto an absorbent surface and tapping to remove any liquid. The pellet was suspended in 200 µl of 8 M Urea in 100 mM Tris pH 8.0. TCEP (tris(2-carboxyethyl)phosphine) was added to final concentration of 10 mM and 2-chloroacetamide was added to final concentration of 40 mM, and the solution was vortexed for 5 minutes. Three volumes of 50 mM Tris pH 8.0 were added to the sample to reduce the final urea concentration to 2 M. Mass spectrometry-grade trypsin was added in a 1:50 ratio of trypsin to protein, and the solution was incubated at 37°C for 12 hours. The solution was then acidified using TFA (trifluoroacetic acid; 0.5% final concentration) and mixed. The sample was desalted using C18-StageTips (Thermo Fisher Scientific). The final peptide concentration of the sample was measured using a BCA protein assay.

### Mass spectrometry data-dependent acquisition (DDA)

1 µg of each sample was analyzed by ultra high pressure liquid chromatography (UPLC) coupled with tandem mass spectrometry (LC-MS/MS) using nanospray ionization. The nanospray ionization experiments were performed using a TimsTOF 2 HT hybrid mass spectrometer (Bruker) interfaced with nanoscale reversed-phase UPLC (Evosep One). The Evosep method 30 SPD (30 samples per day) was used with a 15 cm x 150 µm reverse-phase column packed with 1.5 µm C18-beads (PepSep, Bruker) at 58°C. The analytical columns were connected with a fused silica ID emitter (10 μm ID; Bruker Daltonics) inside a nanoelectrospray ion source (Captive spray source; Bruker). The mobile phases comprised 0.1% formic acid (FA) as solution A and 0.1% FA/99.9% acetonitrile (ACN) as solution B. The mass spectrometry settings for the TimsTOF 2 HT were as follows: Parallel Accumulation–Serial Fragmentation (PASEF) method for standard proteomics. The values for mobility-dependent collision energy ramping were set to 95 eV at an inverse reduced mobility (1/k_0_) of 1.6 V s/cm^2^ and 23 eV at 0.73 V s/cm^2^. Collision energies were linearly interpolated between these two 1/k_0_ values and kept constant above or below. No merging of TIMS scans was performed. Target intensity per individual PASEF precursor was set to 20,000. The scan range was set between 0.6 and 1.6 V s/cm^2^ with a ramp time of 166 ms. Fourteen (14) PASEF MS/MS scans were triggered per cycle (2.57 s) with a maximum of seven precursors per mobilogram. Precursor ions in an m/z range between 100 and 1700 with charge states ≥3+ and ≤8+ were selected for fragmentation. Active exclusion was enabled for 0.4 min (mass width 0.015 Th, 1/k_0_ width 0.015 V s/cm^2^).

### Mass spectrometry data analysis and normalization

Peptide identification and quantification was carried out using Peaks Studio 12.5 (Bioinformatics Solutions Inc.)^63^, using sequence databases for the *E. coli* and phage Goslar proteomes; only peptides assigned to Goslar proteins were further processed. Label-free quantitation (LFQ) peak areas for each peptide were summed to generate total LFQ peak area per protein, then this value was divided by the predicted molecular weight of each protein to generate LFQ area/mass for each protein. The total LFQ area/mass was calculated by summing across all detected Goslar proteins, then each protein’s LFQ area/mass was divided by this sum to generate LFQ copy number fraction for each protein. For structural proteins with a known copy number per virion, the LFQ copy number fraction was divided by this number to generate the LFQ fraction per copy. To estimate the copy number per virion of proteins not detected in our structures, we used the LFQ fraction per copy for three known high-copy proteins: MCP (gp041) at 0.00382% per copy, tail tube (gp217) at 0.00502% per copy, and tail sheath (gp218) at 0.00901% per copy to define an estimated median LFQ fraction per copy of ∼0.006% +/− ∼0.003%.

Other structural proteins like those in the portal vertex complex were not used for this calculation, due to uncertainty about the total copy number of these proteins per virion. The estimated copy number using the mean LFQ fraction per copy of 0.006% is reported in the main text; upper and lower estimates for each protein are available in **Table S4**).

### Infections and Microscopy

#### Plasmid construction and bacterial transformation

Selected Goslar genes were closed as fusions to GFPmut1 at either the N- or C-terminus and expressed from vector pDSW206 (ref. ^28^) modified to include a second RBS, “TTTAAGAAGGAGAAATTCACC”, to drive increased protein expression compared to the standard vector. 1 μL of 20 μg/mL plasmid was added to 30-50 μL of electroporation-competent *E. coli* MC1000 cells (strain EA258; prepared by washing in 10% glycerol) and electroporated with 2.5 kV. After transformants recovered in 1 mL SOC at 37°C for 60 minutes, they were spread on LB plates with 100 μg/mL ampicillin and incubated overnight at 37°C.

#### Fluorescence microscopy

Fluorescence microscopy experiments were performed in biological triplicate. Host cells were suspended in LB at OD600 0.6 as measured by nanodrop. 5 μL were spread on the surface of 1% agarose, 25% LB imaging pads (containing 100 μg/mL ampicillin plus 10 μM or 100 μM isopropyl β-D-1-thiogalactopyranoside (IPTG) for induction of GFP fusion protein expression) in single-well concavity glass slides. After the slides incubated for 1.5-2 hours at 37°C without coverslips in a humidor, 10 μL of ∼10^10^ PFU/mL Goslar lysate were spread onto the imaging pads and the pads were incubated further at 37°C until the desired infection time point. For fluorescence microscopy, pads were stained with 8 μL of dye mix containing 25 μg/mL DAPI and 3.75 μg/mL FM4-64 at room temperature ∼5 minutes before imaging. Cells were imaged with 5-12 slices in the Z-axis at 0.2 or 0.3 μm increments. All microscopy was performed on a DeltaVision Elite Deconvolution microscope (Applied Precision, Issaquah, WA, USA) via DeltaVision SoftWoRx (version 6.5.2). Images were deconvolved in DeltaVision SoftWoRx (version 6.5.2).

### DNA Cloning and Protein Expression

For protein expression, Goslar gp022 (NCBI:YP_009820707.1) was cloned into UC Berkeley Macrolab vector 2-BT (Addgene #29666) to generate an N-terminal TEV protease-cleavable His_6_-tagged construct. Proteins were expressed in *E. coli* Rosetta2 pLysS (EMD Millipore) by growing cells at 37°C to an optical density at 600 nm (OD_600_) of 0.6–0.8, followed by induction with 0.33 mM IPTG. Cultures were incubated overnight at 20°C with shaking. After 14-16 hours, cells were collected by centrifugation, and pellets were resuspended in ice-cold resuspension buffer containing 20 mM Tris-HCl pH 7.5, 300 mM NaCl, 10 mM imidazole, 10% glycerol and 2 mM β-mercaptoethanol. Resuspended cells were lysed by sonication (Branson Sonifier), and the lysate was clarified by centrifugation. The protein was purified using Ni^2+^ affinity chromatography (Ni-NTA Superflow, Qiagen), with one wash in resuspension buffer plus 2 M urea to disrupt large protein aggregates. Protein was eluted with resuspension buffer plus 250 mM imidazole.

### Negative-Stain EM

5 µl purified gp22 protein was applied to a glow discharged 400 mesh Copper formvar/Carbon EM grid (Ted Pella #01754-F) for 5 minutes. The grid was washed 3 times with 0.22 μm-filtered double-distilled water and incubated for 1 minute with 0.22 μm-filtered 2% uranyl acetate (Ladd Research #23620) in double-distilled water. The grid was blotted for 3 seconds using Whatman Grade 1 filter paper (Whatman #1001-085) and air-dried on inverted tweezers. Images were captured with a FEI Tecnai Spirit G2 BioTWIN operated at 80 KeV and equipped with a bottom-mounted 4kx4k FEI Eagle camera.

## DATA AVAILABILITY

Final molecular models and refined cryoEM maps are available at the RCSB Protein Data Bank (http://rcsb.org) and Electron Microscopy Data Bank (https://www.ebi.ac.uk/emdb/) under the following accession codes: Full Goslar virion (EMD-76130); Empty capsid (PDB_000011ML, EMD-75837); Filled capsid (PDB_000011MK, EMD-75836); Capsid C5 penton (PDB_000011FG, EMD-75663); Capsid C3 hexon (PDB_000011BT, EMD-75608); Portal and upper neck (PDB_000011ED, EMD-75646); Lower neck (PDB_000011LC, EMD-75794); Tail (PDB_000011HF, EMD-75688); Tail tube (PDB_000011EC, EMD-75645); Tail sheath (PDB_000011EI, EMD-75649); Baseplate (PDB_000011LX, EMD-75829); Peripheral baseplate (PDB_000011PB, EMD-75910).

## ACKNOWLEDGEMENTS

The authors thank members of the Corbett and Pogliano labs, along with J. Meyer, E. Villa, and D. Pride for helpful conversations. The authors acknowledge support from the Howard Hughes Medical Institute Emerging Pathogens Initiative (to J.P. and K.D.C) and the National Institutes of Health (R01 GM129245 to J.P., R35 GM144121 to K.D.C.). The authors acknowledge the facilities of the UC San Diego cryo-EM facility, part of the Goeddel Family Technology Sandbox, along with the scientific and technical assistance of facility staff members Dr. Mariusz Matyszewski and Dr. Inga Kuschnerus. The authors thank the University of California, San Diego - Cellular and Molecular Medicine Electron Microscopy Core (UCSD-CMM-EM Core, RRID: SCR_022039) for providing access to equipment and technical assistance from Guillame Castillon.

## AUTHOR CONTRIBUTIONS

D.B. performed Goslar phage purifications, cryoEM data collection, and cryoEM processing, prepared figures, and wrote the manuscript. Y.G. performed cryoEM data processing. Y.L. and T.F. performed phage infections and fluorescence imaging. M.G. performed mass spectrometry. J.P. obtained funding, interpreted data, and edited the manuscript. K.D.C. obtained funding, oversaw the project, prepared figures, and wrote the manuscript.

## COMPETING INTERESTS

J.P. has an equity interest in Linnaeus Bioscience Incorporated and receives income. The terms of this arrangement have been reviewed and approved by the University of California, San Diego in accordance with its conflict-of-interest policies. The other authors declare no competing interests.

## SUPPLEMENTAL INFORMATION

**Table S1. CryoEM data collection and structure determination of intact Goslar virions**

*See attached Excel workbook*

**Table S2. CryoEM data collection and structure determination of Goslar substructures**

*See attached Excel workbook*

**Table S3.**
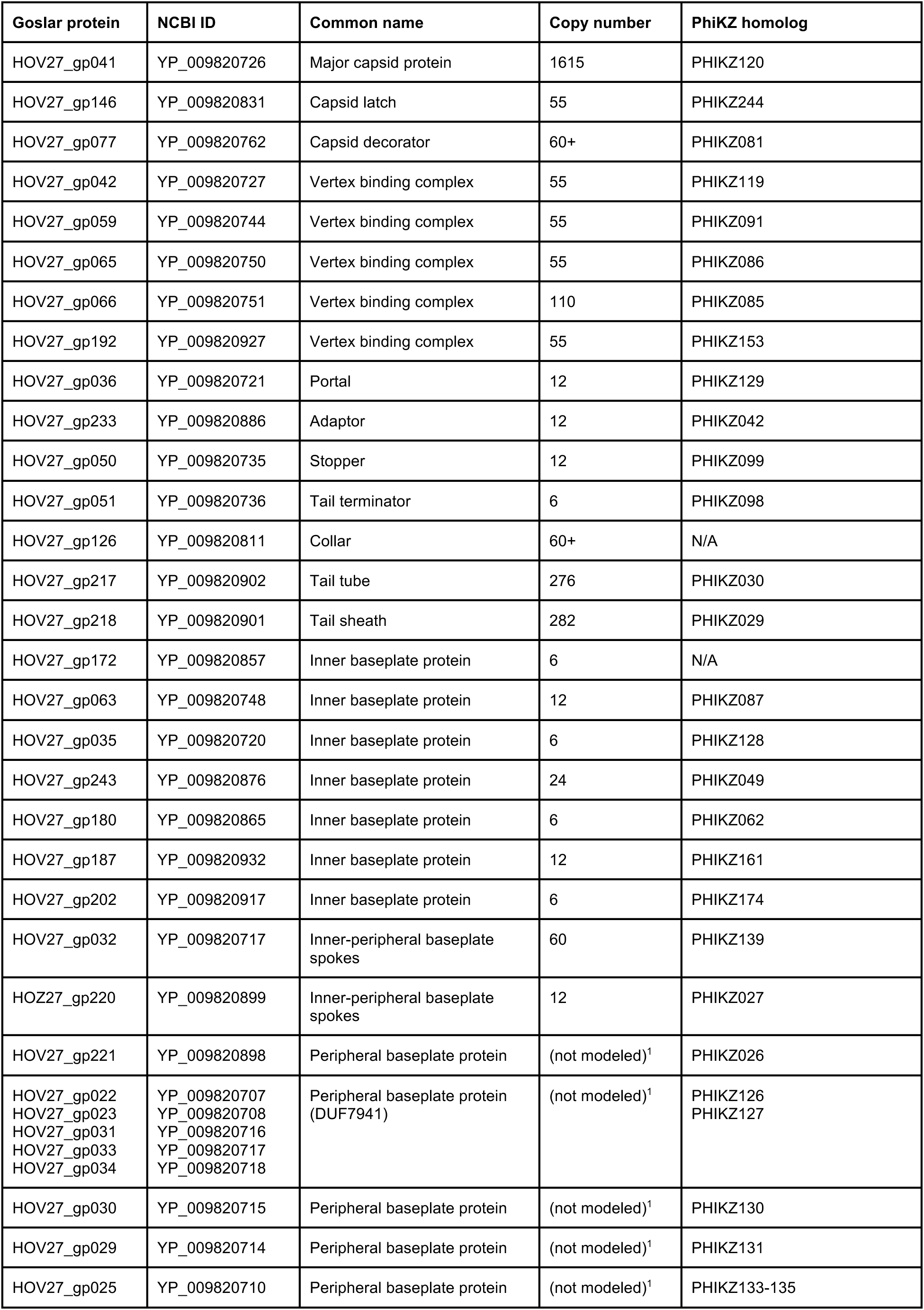
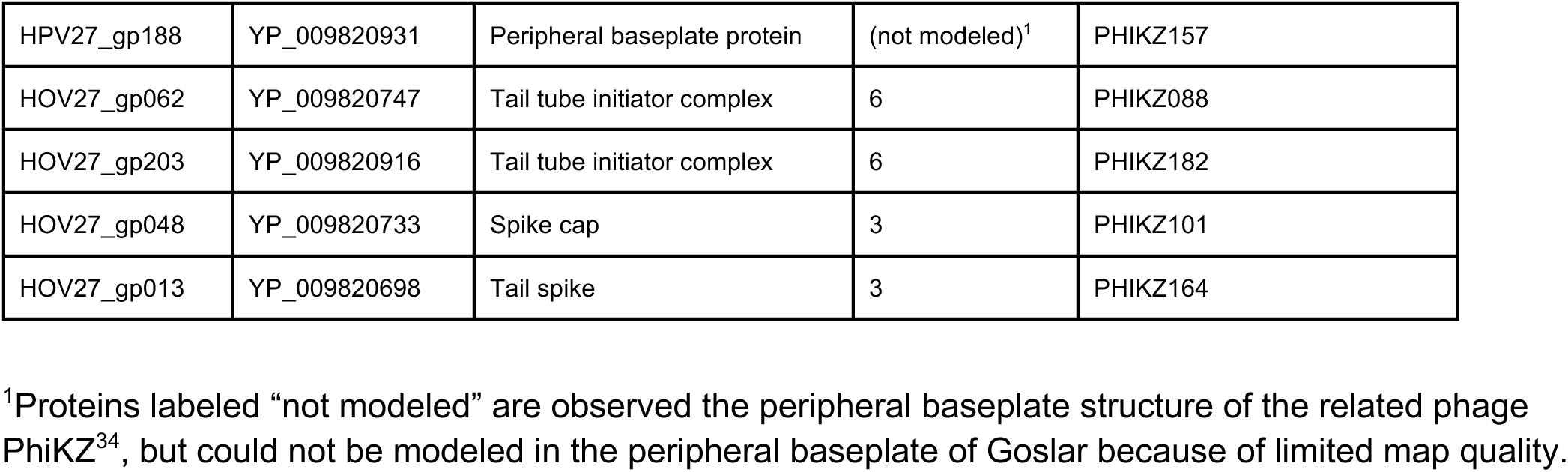
Goslar structural proteins.

**Table S4. LFQ mass spectrometry of purified Goslar virions**

*See attached Excel workbook*

**Figure S1.**
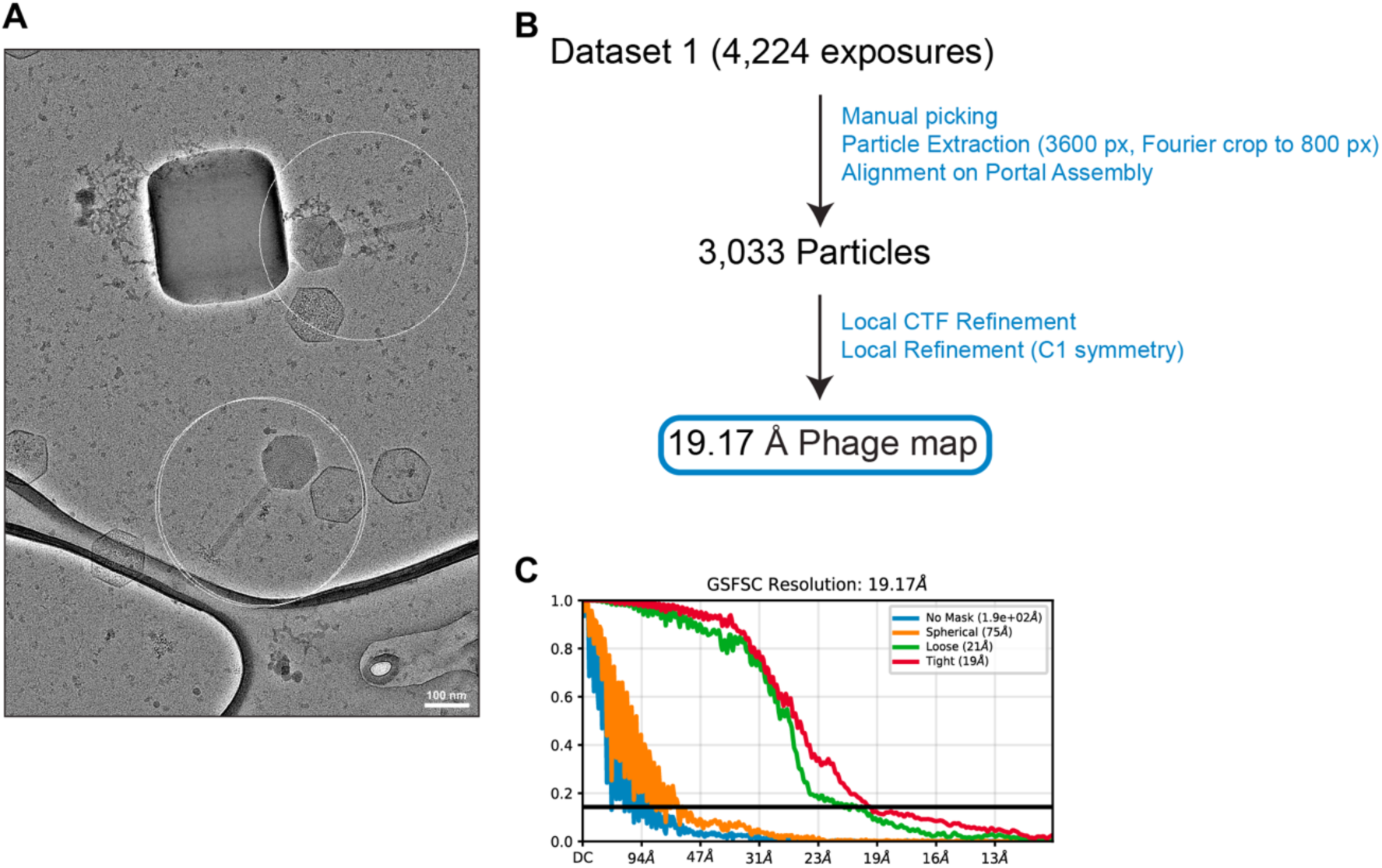
CryoEM workflow for asymmetric reconstruction of Goslar phage. A. Example cryoEM micrograph from dataset 1 (collected in super-resolution mode with pixel size of 1.3 Å), with white circles indicating manually-picked intact Goslar phages. Also visible are several empty capsids without tails (see **Figure S4**). Scale bar: 100 nm. B. CryoEM workflow for structure determination of intact Goslar phages. See **Table S1** for data collection and refinement details. C. Gold-standard Fourier Shell Correlation (GSFSC) curves for the 19.17 Å resolution icosahedral map.

**Figure S2.**
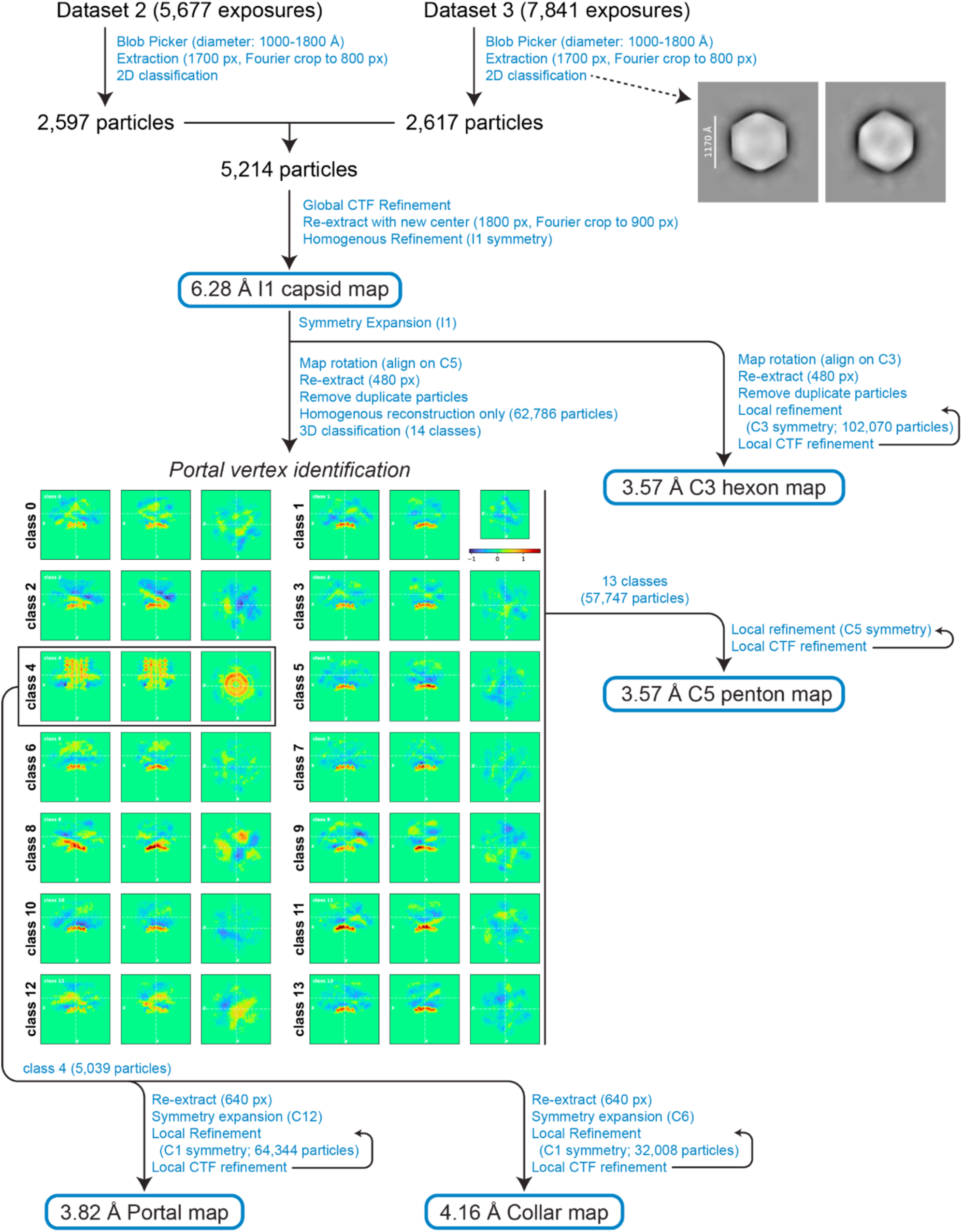
CryoEM workflow for Goslar capsid, portal, and neck reconstructions. CryoEM workflow for structure determination of Goslar capsid, portal, and neck. From two datasets, capsid particles were picked using the blob picker, extracted, then combined for an initial 3D reconstruction with icosahedral symmetry. Particles were then symmetry-expanded, maps were rotated to align on either the five-fold (C5) or three-fold (C3) symmetry axis of the capsid, then re-extracted. For particles aligned on the C3 axis, iterative local refinement and local CTF estimation yielded a 3.57 Å resolution C3 “hexon” map. For particles aligned on the C5 axis, homogeneous reconstruction and 3D classification identified the vertex of each particle associated with the portal. These particles were subjected to iterative local refinement and local CTF estimation with C12 symmetry to generate a 3.82 Å resolution portal/upper neck map, or with C6 symmetry to generate a 4.16 Å resolution lower neck map. All vertices not associated with a portal were combined and refined with C5 symmetry to generate a 3.57 Å resolution C5 “penton” map. See **Table S2** for data collection and refinement details.

**Figure S3.**
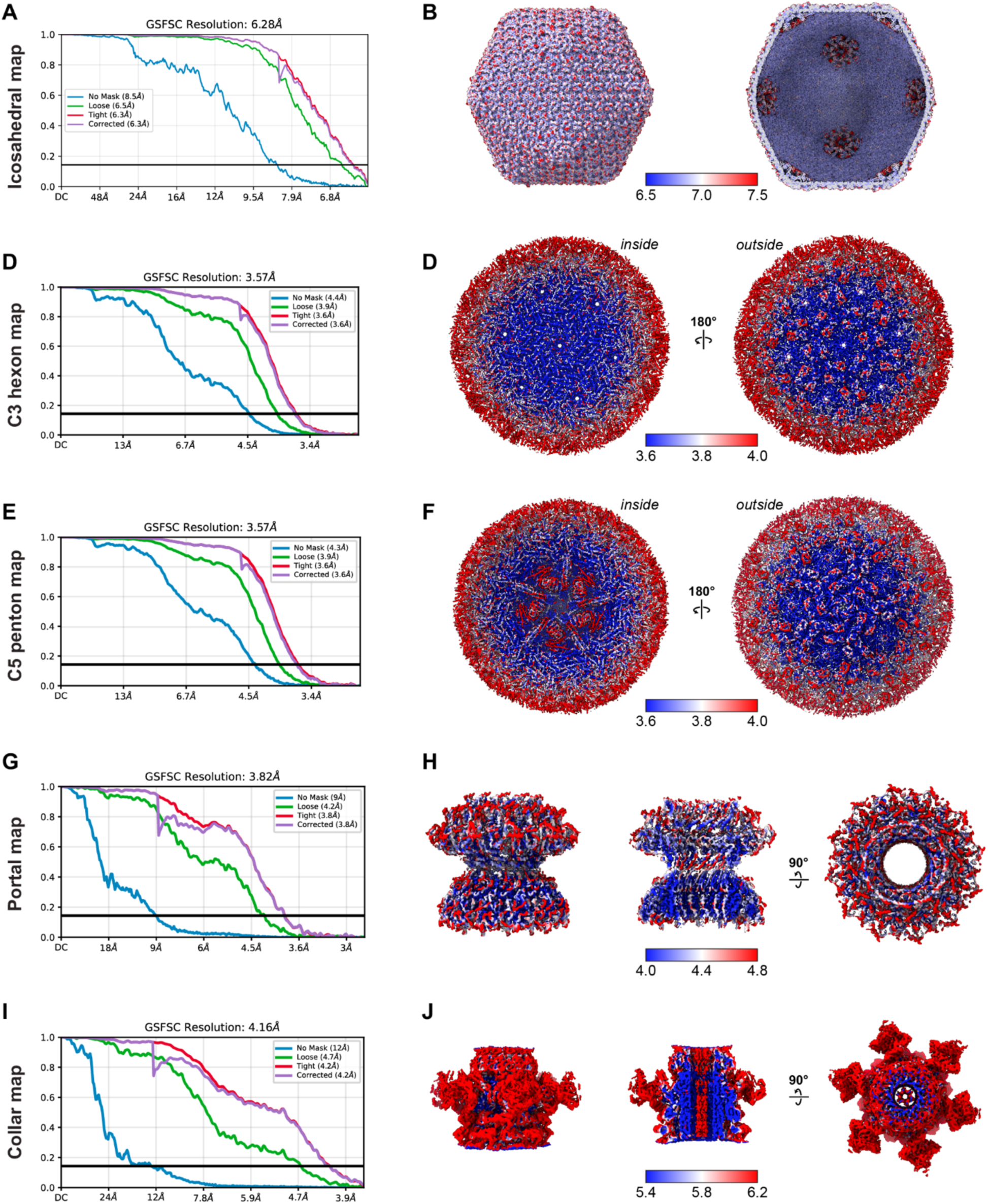
Validation of Goslar capsid, portal, and neck reconstructions. A. Gold-standard Fourier Shell Correlation (GSFSC) curves for the 6.28 Å resolution icosahedral map. B. Exterior (left) and cutaway (right) views of the icosahedral map, colored by local resolution. C. GSFSC curves for the 3.57 Å resolution C3 “hexon” map. D. Inside (left) and outside (right) views of the C3 “hexon” map, colored by local resolution. E. GSFSC curves for the 3.57 Å resolution C5 “penton” map. F. Inside (left) and outside (right) views of the C5 “penton” map, colored by local resolution. G. GSFSC curves for the 3.82 Å resolution C12 portal map. H. Side view (left), cutaway view (center), and top view (right) of the C12 portal/upper neck map, colored by local resolution. I. GSFSC curves for the 4.16 Å resolution C6 lower neck map. J. Side view (left), cutaway view (center), and top view (right) of the C6 lower neck map, colored by local resolution.

**Figure S4.**
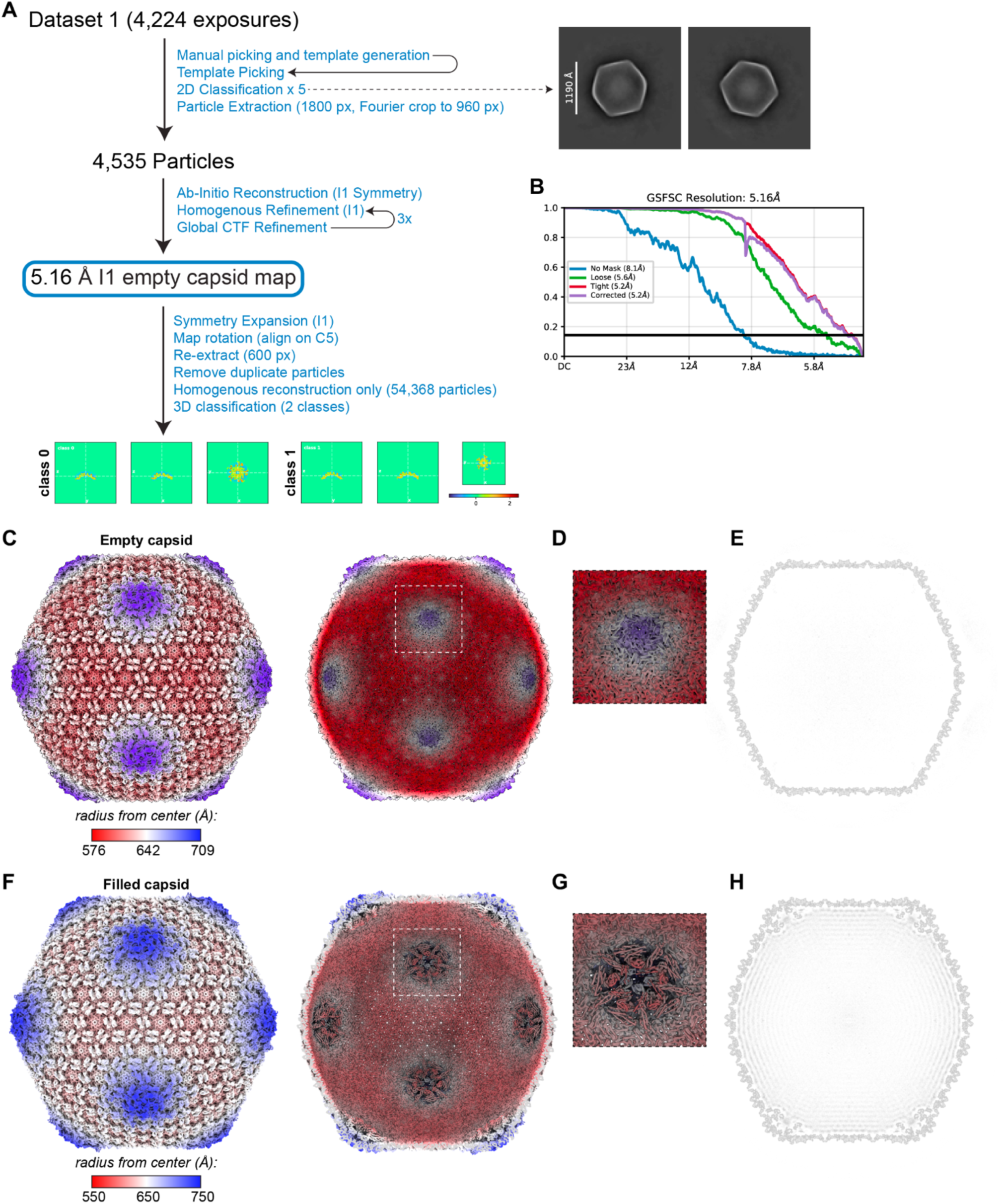
CryoEM workflow for empty capsid reconstruction. A. CryoEM workflow for structure determination of Goslar empty capsids. 3D classification (results of a trial with 2 classes is shown; trials with up to 10 classes showed consistent results) did not reveal a portal vertex. See **Table S1** for data collection and refinement details. B. Gold-standard Fourier Shell Correlation (GSFSC) curves for the 5.16 Å resolution icosahedral map. C. Exterior and cutaway views of the icosahedral map for empty capsids, colored by radius from the center. D. Closeup view of the inside of one C5 vertex in the empty capsid, showing the lack of a vertex binding complex. E. Cross-section of the icosahedral map for empty capsids. F. Exterior and cutaway views of the icosahedral map for DNA-filled capsids (see **Figures S2** and **S3**), colored by radius from the center. G. Closeup view of the inside of one C5 vertex in the DNA-filled capsid, showing the vertex binding complex. H. Cross-section of the icosahedral map for the DNA-filled capsid, showing the vertex-binding complex and layers of density corresponding to DNA.

**Figure S5.**
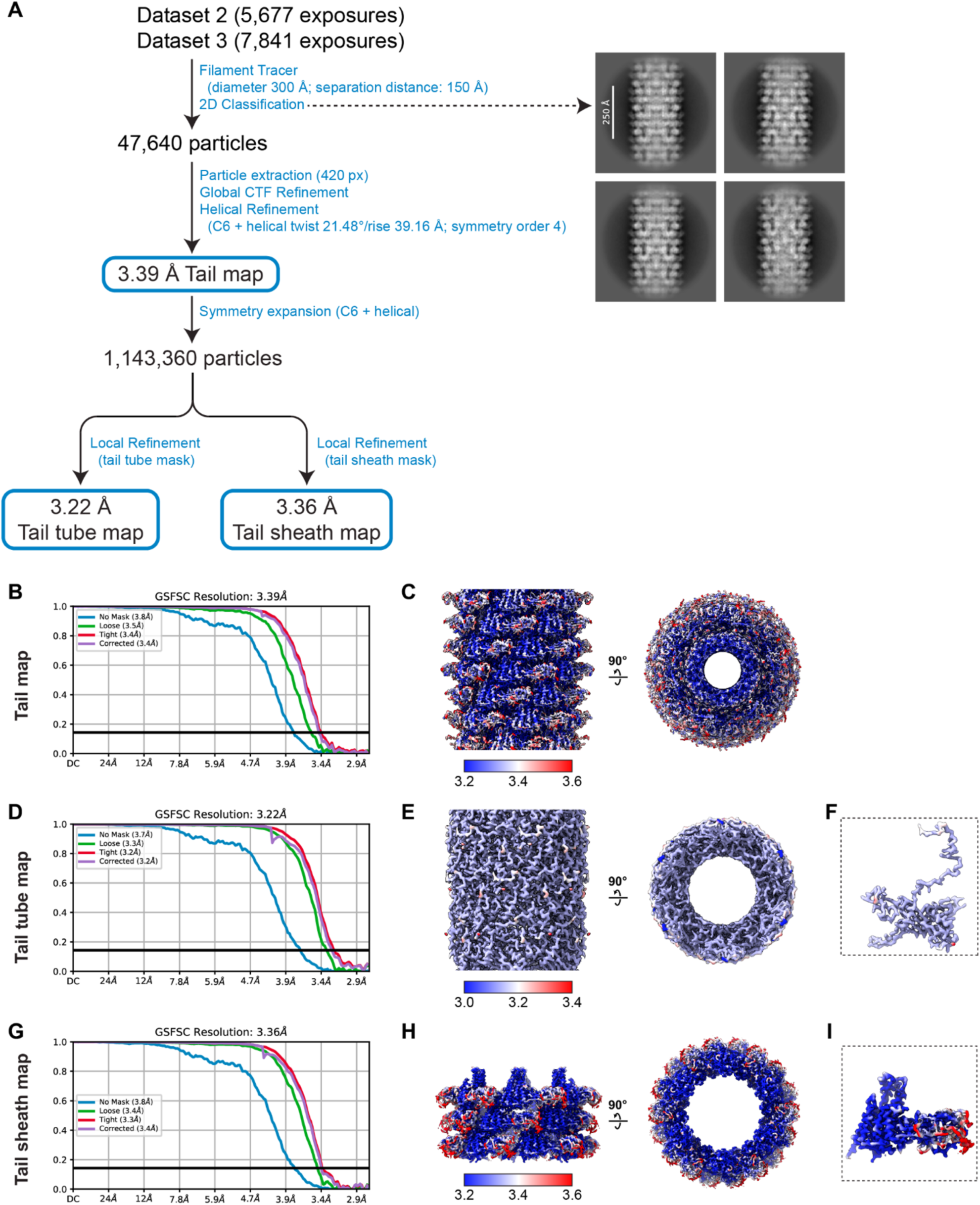
CryoEM workflow for Goslar tail reconstructions. A. From two datasets, the cryoSPARC Filament Tracer was used to pick particles, followed by 2D classification to yield a 47,640 particle set. Helical refinement with C6 symmetry yielded a 3.39 Å resolution tail map. Symmetry expansion and local refinement with masks yielded a 3.22 Å resolution tail tube map, and a 3.36 Å resolution tail sheath map. See **Table S2** for data collection and refinement details. B. GSFSC curves for the 3.39 Å resolution tail map. C. Side and top views of the 3.39 Å resolution tail map, colored by local resolution. D. GSFSC curves for the 3.22 Å resolution tail tube map. E. Side and top views of the 3.22 Å resolution tail map, colored by local resolution. F. View of a single tail tube protein protomer, colored by local resolution. G. GSFSC curves for the 3.36 Å resolution tail sheath map. H. Side and top views of the 3.36 Å resolution tail sheath map, colored by local resolution. I. View of a single tail sheath protein protomer, colored by local resolution.

**Figure S6.**
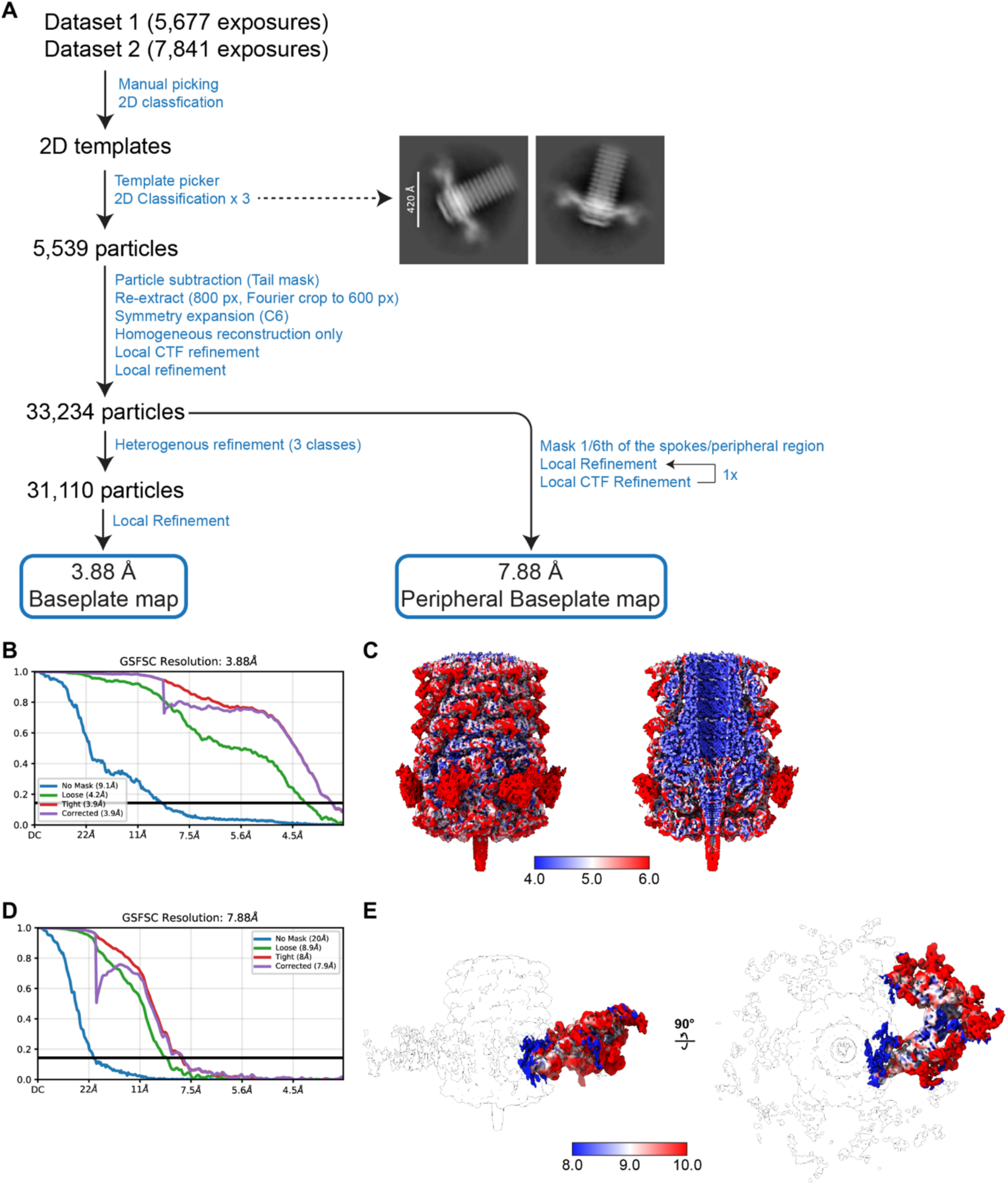
CryoEM workflow for Goslar baseplate reconstruction. A. From two datasets, manually-picked particles were used to generate templates for template-based particle picking. After 2D classification, a 5,539 particle set was used for initial reconstruction and refinement with C6 symmetry. Symmetry expansion followed by local refinement resulted in a final map at 3.88 Å resolution. To better resolve the spokes and peripheral region, a manual mask around 1/6th of that region was used for a second local refinement, resulting in a 7.88 Å map. See **Table S2** for data collection and refinement details. B. GSFSC curves for the 3.88 Å resolution baseplate map. C. Side and cutaway views of the 3.88 Å resolution baseplate map, colored by local resolution. D. GSFSC curves for the 7.88 Å resolution spoke/peripheral baseplate map. E. Side and top views of the 7.88 Å resolution spoke/peripheral baseplate map, colored by local resolution.

**Figure S7.**
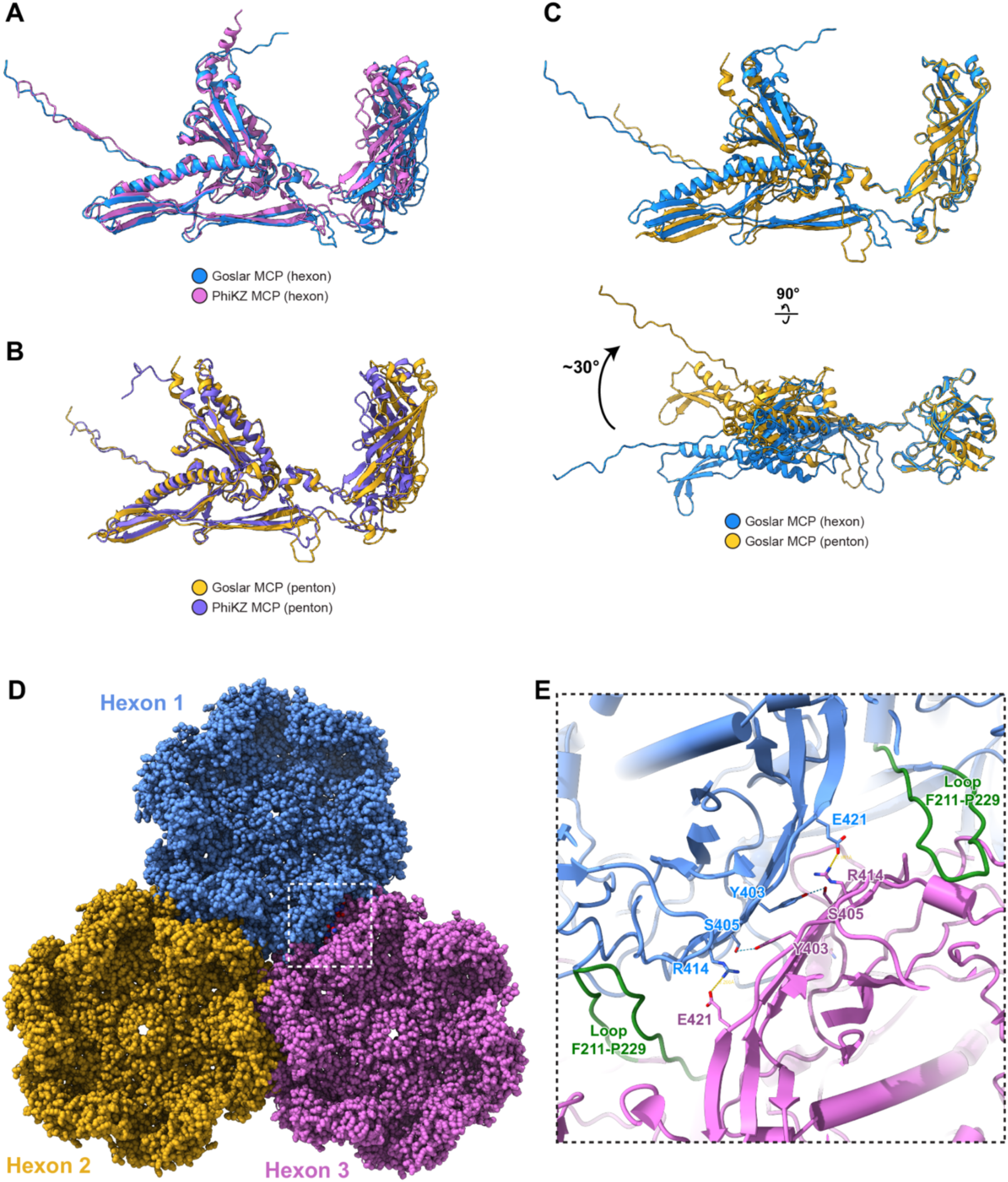
Structure and packing of the Goslar major capsid protein. A. Overlay of the Goslar major capsid protein in the hexon (gp41; blue) with that of PhiKZ (PDB ID 8Y6V)^41^ (purple). B. Overlay of Goslar gp41 in the penton (yellow) with that of PhiKZ. C. Two views of Goslar gp41 in the hexon (blue) and penton (yellow), showing rotation of the I domain relative to the rest of the protein. D. View of three gp41 hexons packing against one another in the capsid. E. Closeup view of the interactions between gp41 protomers in neighboring hexons.

**Figure S8.**
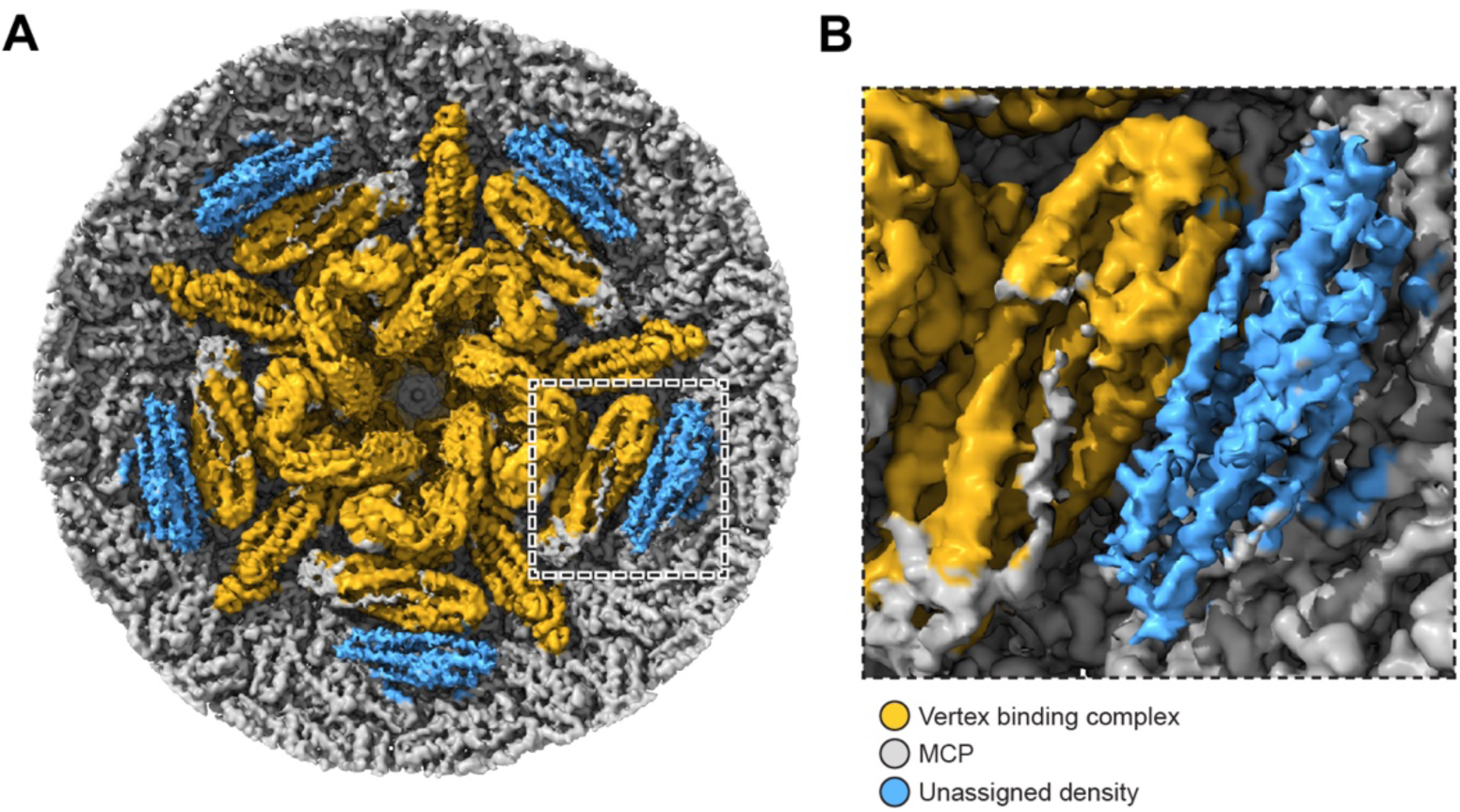
Unassigned density near the vertex binding complex. A. CryoEM density for the vertex binding complex at low contour level, viewed from inside the Goslar capsid. Density corresponding to the confidently-built vertex binding complex is colored yellow; density corresponding to the capsid (major capsid protein; MCP) is gray), and unassigned density is blue. B. Closeup view of one unassigned density (blue). The neighboring yellow density corresponds to gp65.

**Figure S9.**
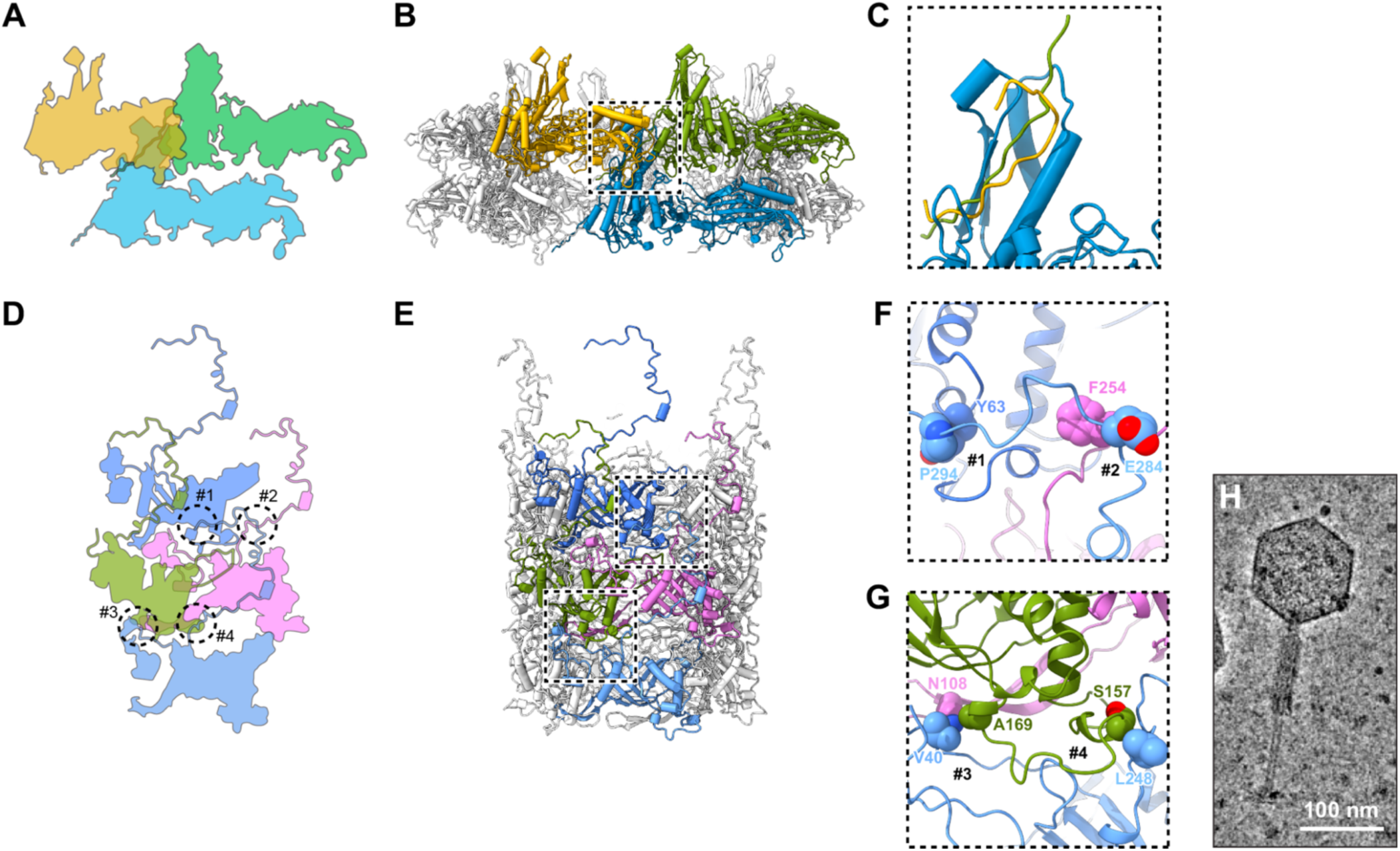
Goslar tail architecture. A. Outline schematic showing the positioning of three interacting protomers of the tail sheath protein gp218. B. Cartoon view of three interacting tail sheath protomers in the context of the full tail sheath assembly (two layers shown). C. Closeup of the critical interaction between the C-terminal domain of one tail sheath protomer (blue) with two other protomers (yellow and green) in the layer above. D. Outline schematic showing the positioning of four interacting protomers of the tail tube protein gp217. Key interactions are marked #1 through #4. E. Cartoon view of the four interacting tail tube protomers in the contect of the full tail tube assembly (three layers shown). F. Closeup of interactions #1 and #2. G. Closeup of interactions #3 and #4. H. Electron micrograph of a Goslar virion with a contracted tail sheath.

**Figure S10.**
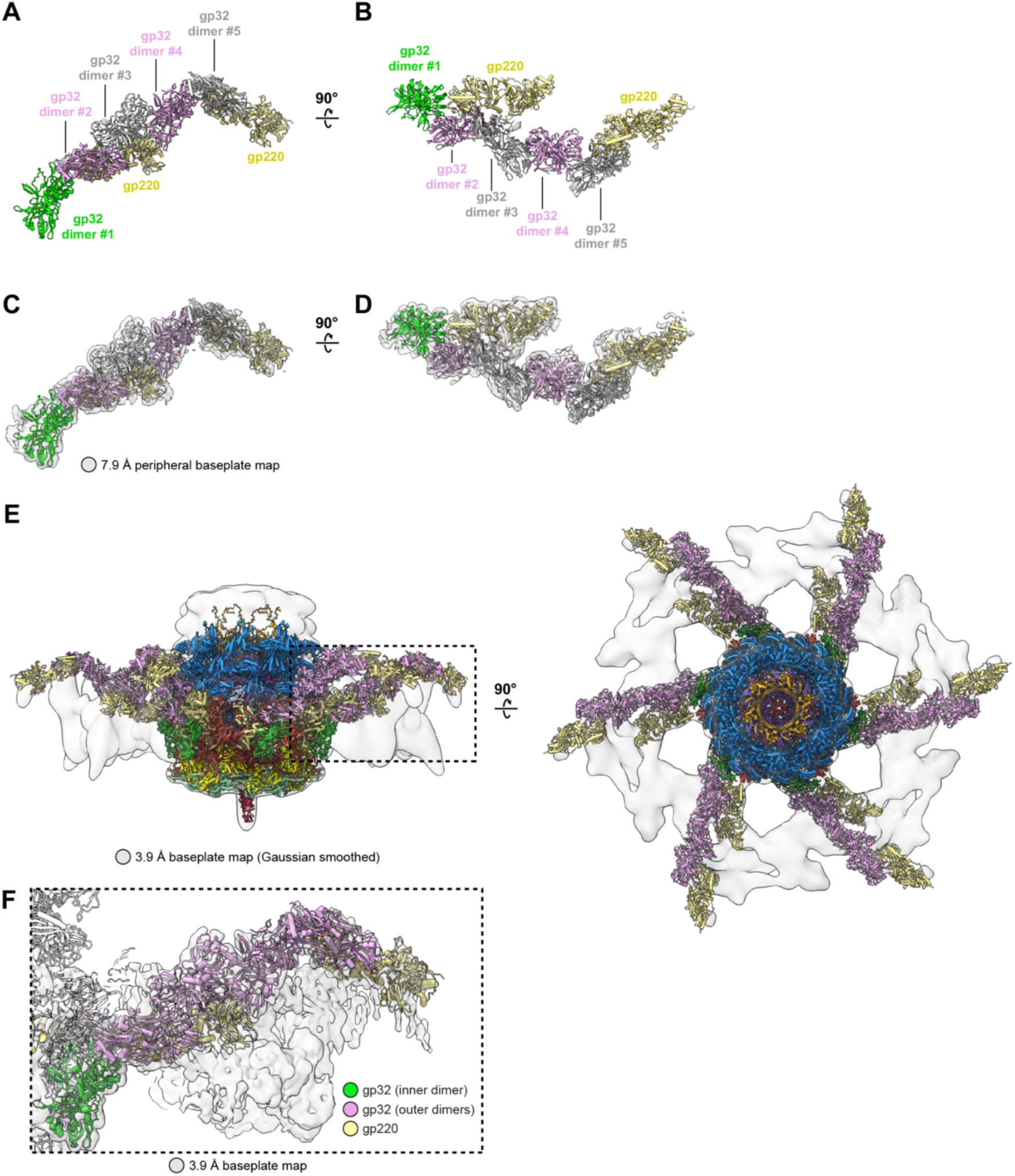
Spoke and peripheral baseplate architecture. A. Side view of one spoke, with five dimers of gp32 and two copies of gp220 labeled. B. Top-down view of one spoke. C. Side view of one spoke, with cryoEM density from the 7.9 Å resolution peripheral baseplate map shown. D. Top-down view of one spoke, with cryoEM density from the 7.9 Å resolution peripheral baseplate map shown. E. Side and top-down views of the entire baseplate molecular model, with atomic models colored by protein as in Figure 6B-D, and Gaussian-smoothed cryoEM density from the 3.9 Å resolution baseplate map shown in gray. F. Closeup of one spoke, with five dimers of gp32 and two copies of gp220 shown as cartoons, and cryoEM density from the 3.9 Å resolution baseplate map (not smoothed) shown in gray.

**Figure S11.**
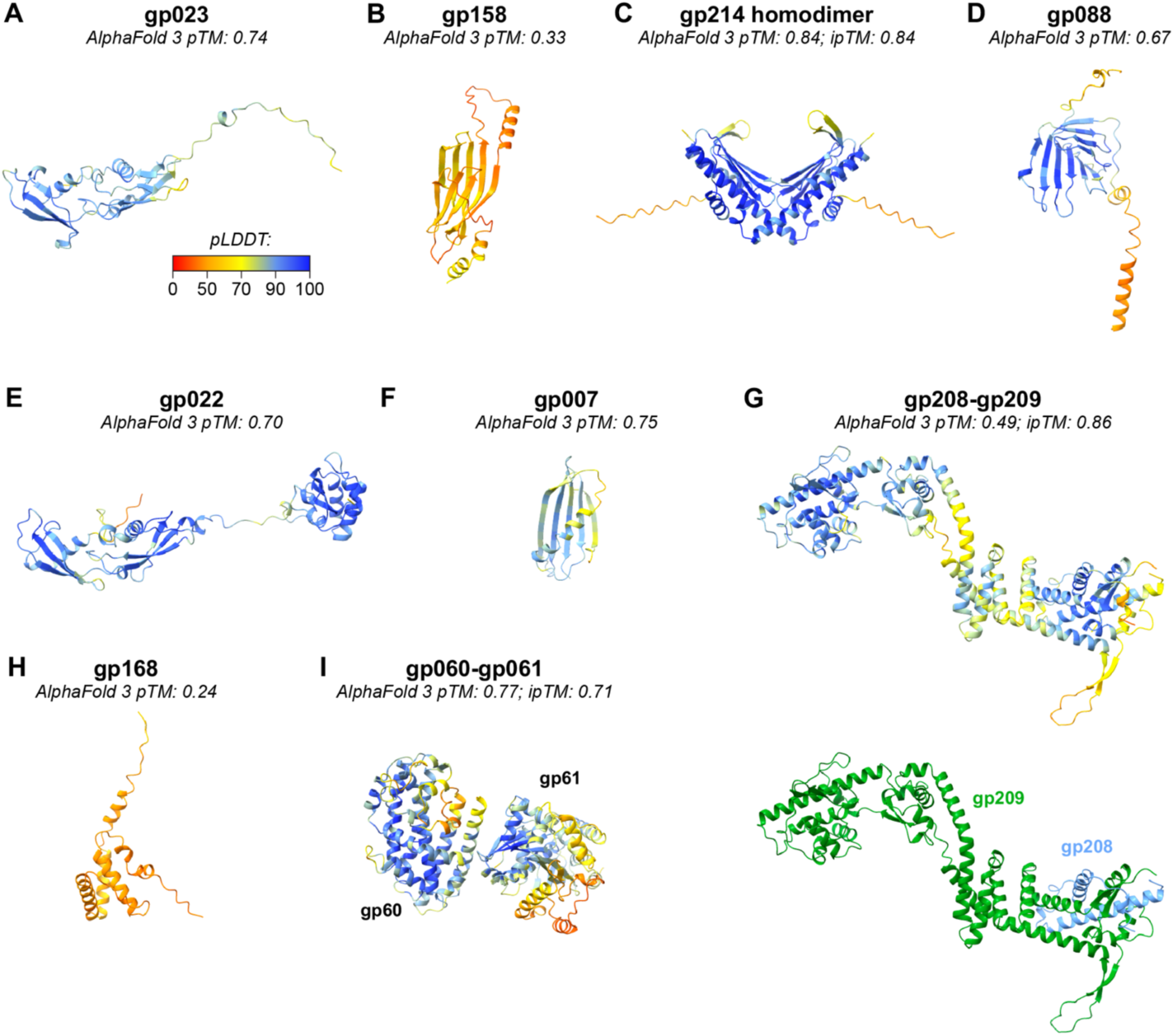
AlphaFold-predicted structures of high-copy Goslar virion proteins. A. Alphafold 3 prediction of Goslar gp023, colored by local confidence (pLDDT). B. Alphafold 3 prediction of Goslar gp158. C. Alphafold 3 prediction of a homodimer of Goslar gp214. D. Alphafold 3 prediction of Goslar gp088. E. Alphafold 3 prediction of Goslar gp022. F. Alphafold 3 prediction of Goslar gp007. G. Alphafold 3 prediction of a heterodimeric Goslar gp208-gp209 complex. Top: colored by pLDDT. Bottom: colored by protein with gp208 blue and gp209 green. H. Alphafold 3 prediction of Goslar gp168. I. Alphafold 3 prediction of a heterodimeric Goslar gp060-gp061 complex, colored by pLDDT.

**Figure S12.**
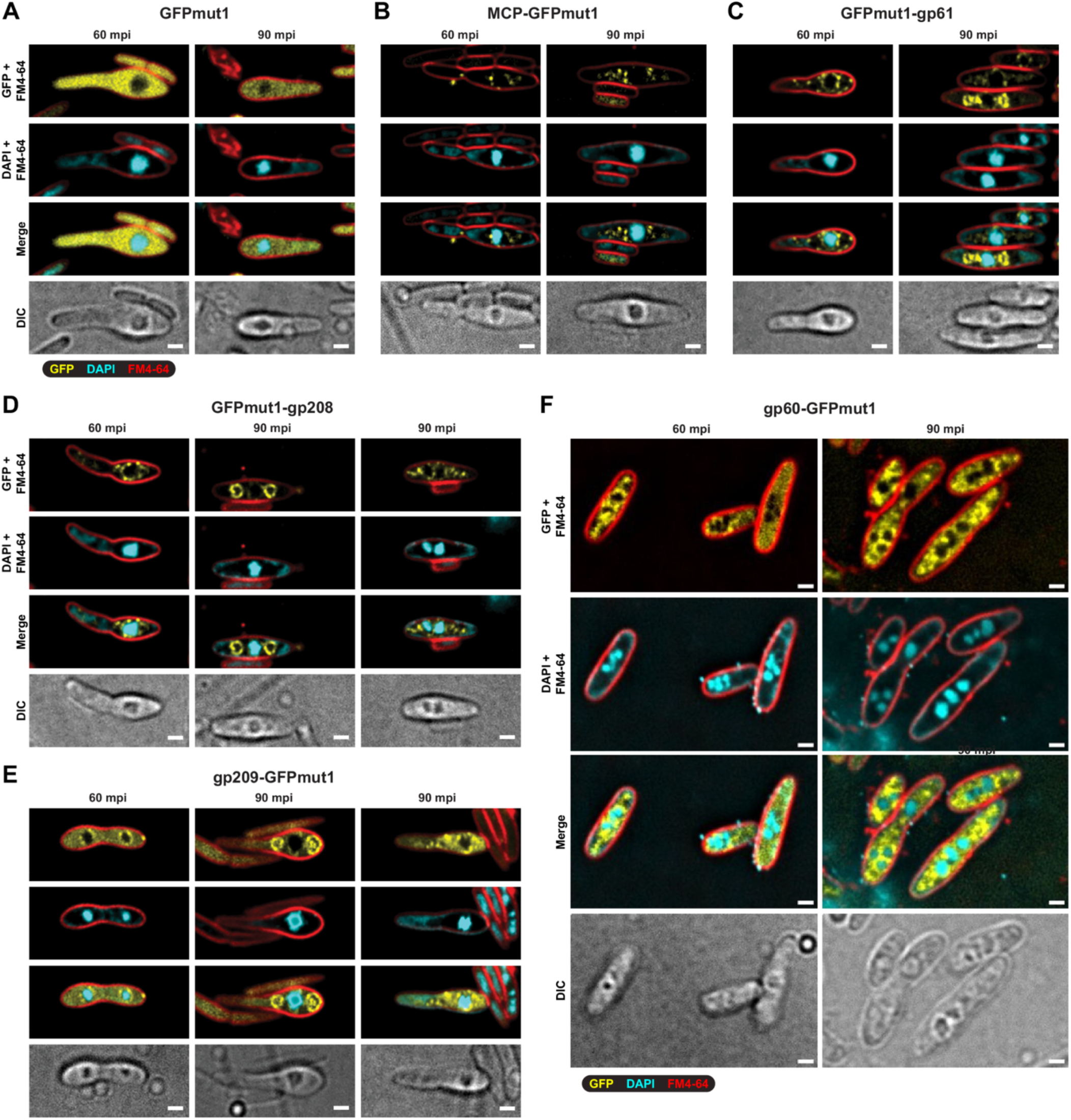
Localization of high-copy virion-associated proteins in Goslar-infected cells. A. Subcellular localization of GFPmut1 in Goslar-infected *E. coli* MC1000 cells, 60 or 90 minutes post infection (mpi). GFP is shown in yellow; DAPI (DNA) in cyan; and FM4-64 (membranes) in red. Differential interference contrast (DIC) imaging is shown at bottom. Scale bar = 1 µm. B. Localization of Goslar gp41 (major capsid protein; MCP) tagged with GFPmut1 at the C-terminus. C. Localization of Goslar gp61 tagged with GFPmut1 at the N-terminus. D. Localization of Goslar gp208 tagged with GFPmut1 at the N-terminus. Two cells are shown at 90 mpi. E. Localization of Goslar gp209 tagged with GFPmut1 at the C-terminus. The cell shown at 60 mpi is double-infected, and shows two phage nuclei. Two cells are shown at 90 mpi. F. Localization of Goslar gp60 tagged with GFPmut1 at the C-terminus.

**Figure S13.**
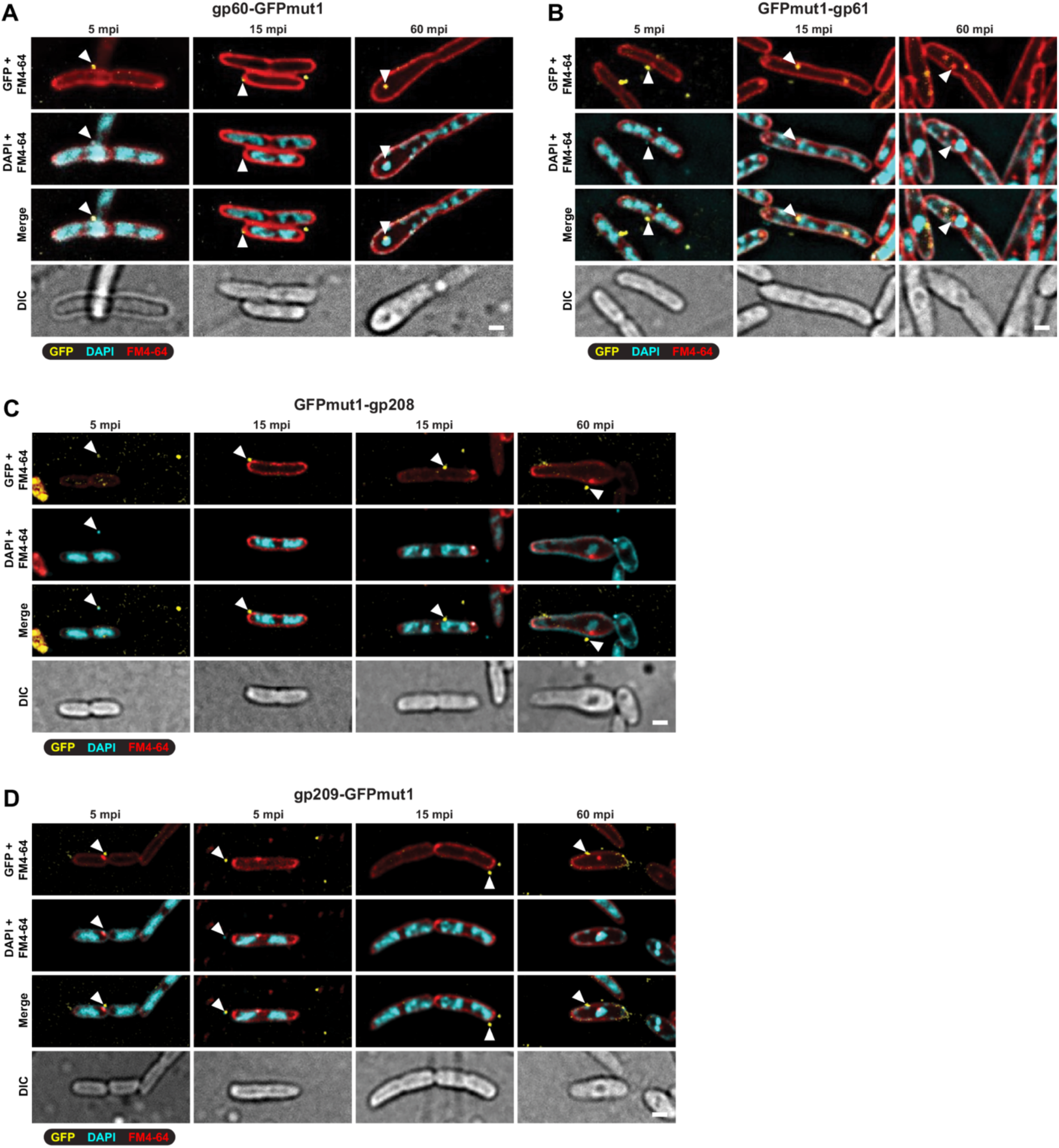
Virion packaging and host injection of Goslar virion-associated proteins. A. Localization of gp60-GFPmut1 packaged into virions and infecting *E. coli* MC1000 cells, at 5, 15, or 60 minutes post infection (mpi). GFP is shown in yellow; DAPI (DNA) in cyan; and FM4-64 (membranes) in red. Differential interference contrast (DIC) imaging is shown at bottom. Scale bar = 1 µm. B. Localization of GFPmut1-gp61 packaged into virions. C. Localization of GFPmut1-gp208 packaged into virions. D. Localization of gp209-GFPmut1 packaged into virions.

**Figure S14.**
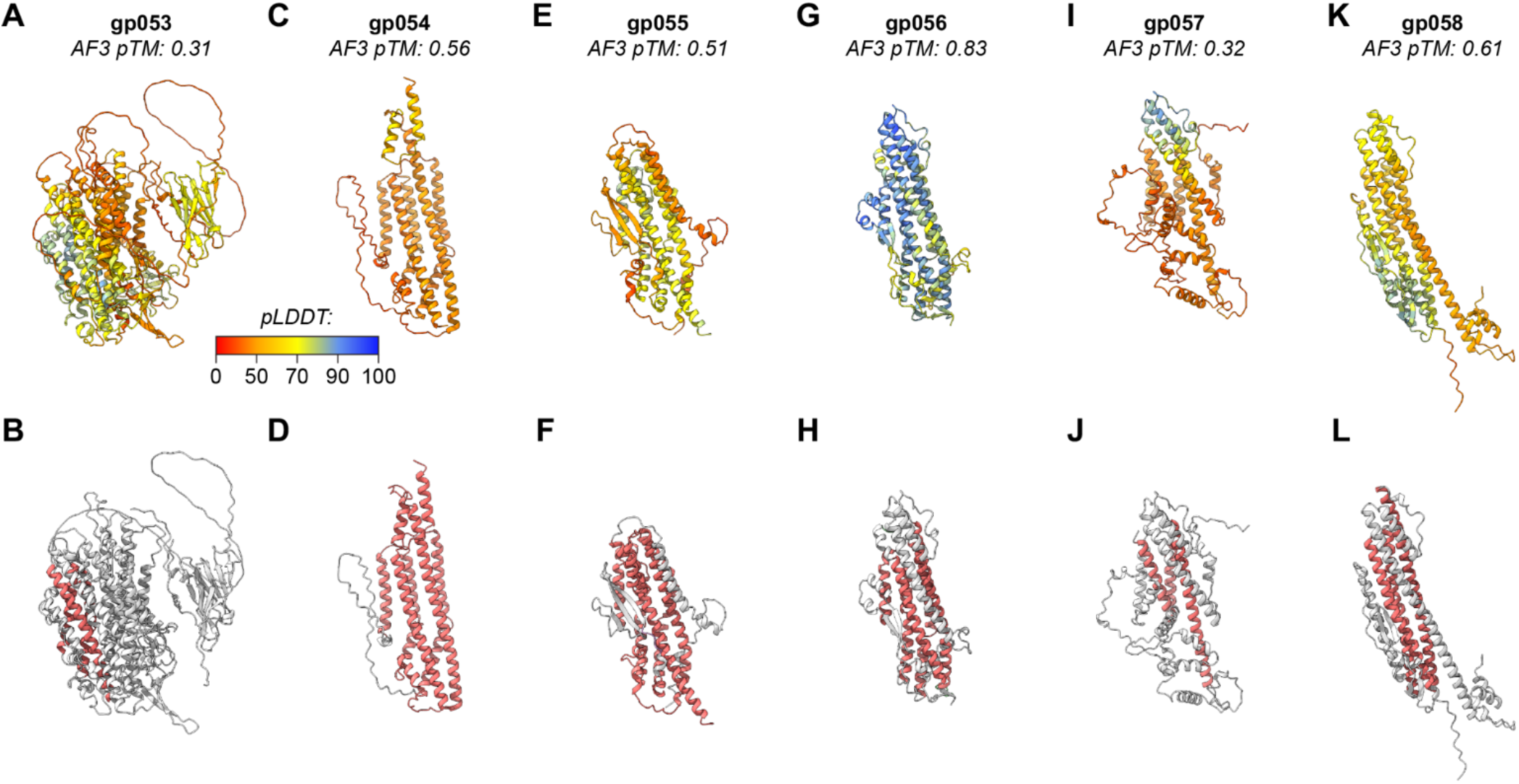
Predicted structures of Goslar PF12699 proteins. A. Alphafold 3 prediction of Goslar gp053, colored by local confidence (pLDDT). B. Alphafold 3 prediction of Goslar gp053, with secondary structure elements corresponding to the core PF12699 fold shown in red. C. Alphafold 3 prediction of Goslar gp054, colored by local confidence (pLDDT). D. Alphafold 3 prediction of Goslar gp054, with secondary structure elements corresponding to the core PF12699 fold shown in red. E. Alphafold 3 prediction of Goslar gp055, colored by local confidence (pLDDT). F. Alphafold 3 prediction of Goslar gp055, with secondary structure elements corresponding to the core PF12699 fold shown in red. G. Alphafold 3 prediction of Goslar gp056, colored by local confidence (pLDDT). H. Alphafold 3 prediction of Goslar gp056, with secondary structure elements corresponding to the core PF12699 fold shown in red. I. Alphafold 3 prediction of Goslar gp057, colored by local confidence (pLDDT). J. Alphafold 3 prediction of Goslar gp057, with secondary structure elements corresponding to the core PF12699 fold shown in red. K. Alphafold 3 prediction of Goslar gp058, colored by local confidence (pLDDT). L. Alphafold 3 prediction of Goslar gp058, with secondary structure elements corresponding to the core PF12699 fold shown in red.

**Figure S15.**
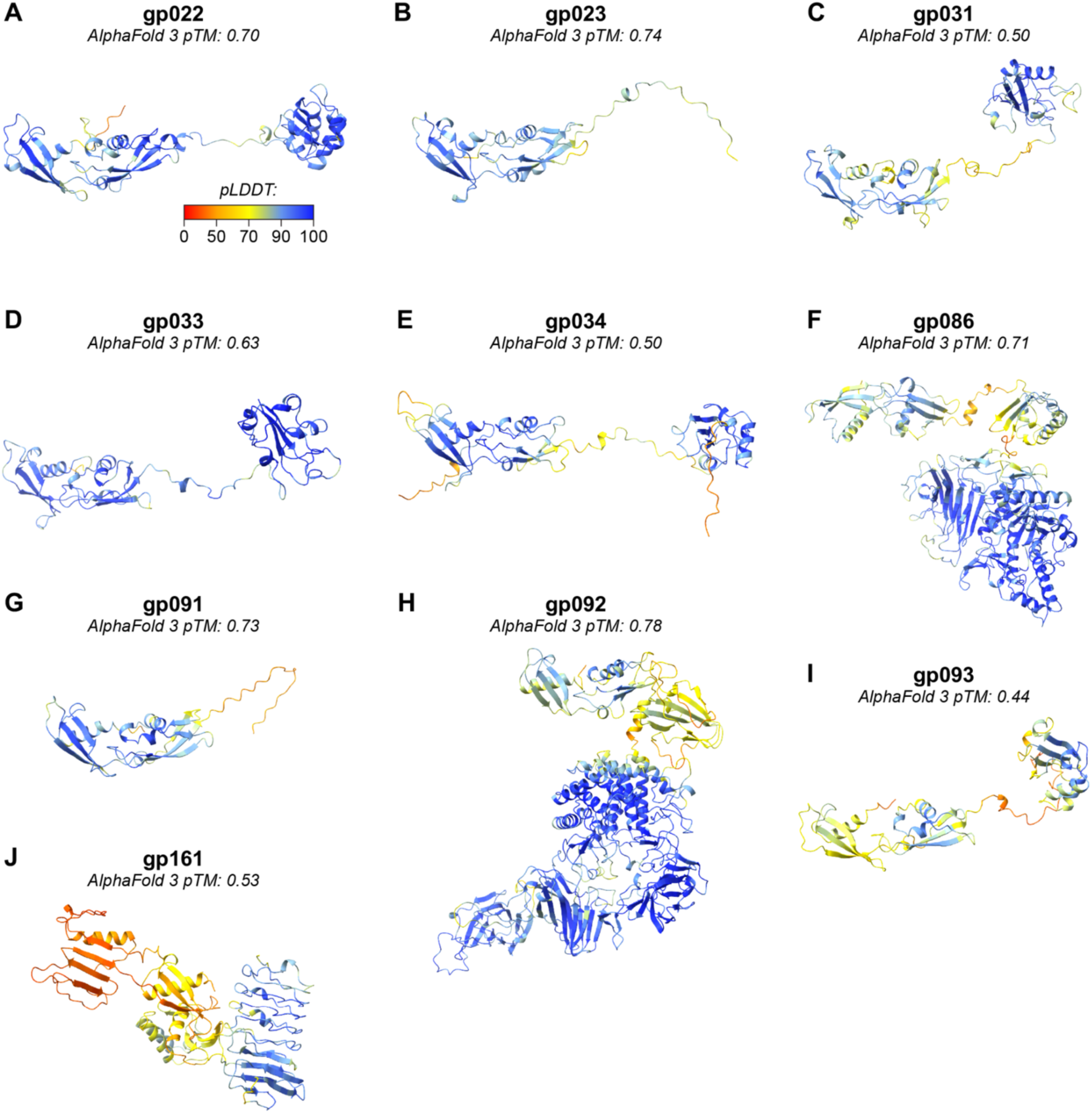
Predicted structures of Goslar DUF7941 proteins. A. AlphaFold 3 prediction of Goslar gp22, colored by local confidence (pLDDT). B. AlphaFold 3 prediction of Goslar gp23. C. AlphaFold 3 prediction of Goslar gp31. D. AlphaFold 3 prediction of Goslar gp33. E. AlphaFold 3 prediction of Goslar gp34. F. AlphaFold 3 prediction of Goslar gp86. G. AlphaFold 3 prediction of Goslar gp91. H. AlphaFold 3 prediction of Goslar gp92. I. AlphaFold 3 prediction of Goslar gp93. J. AlphaFold 3 prediction of Goslar gp161.

**Figure S16.**
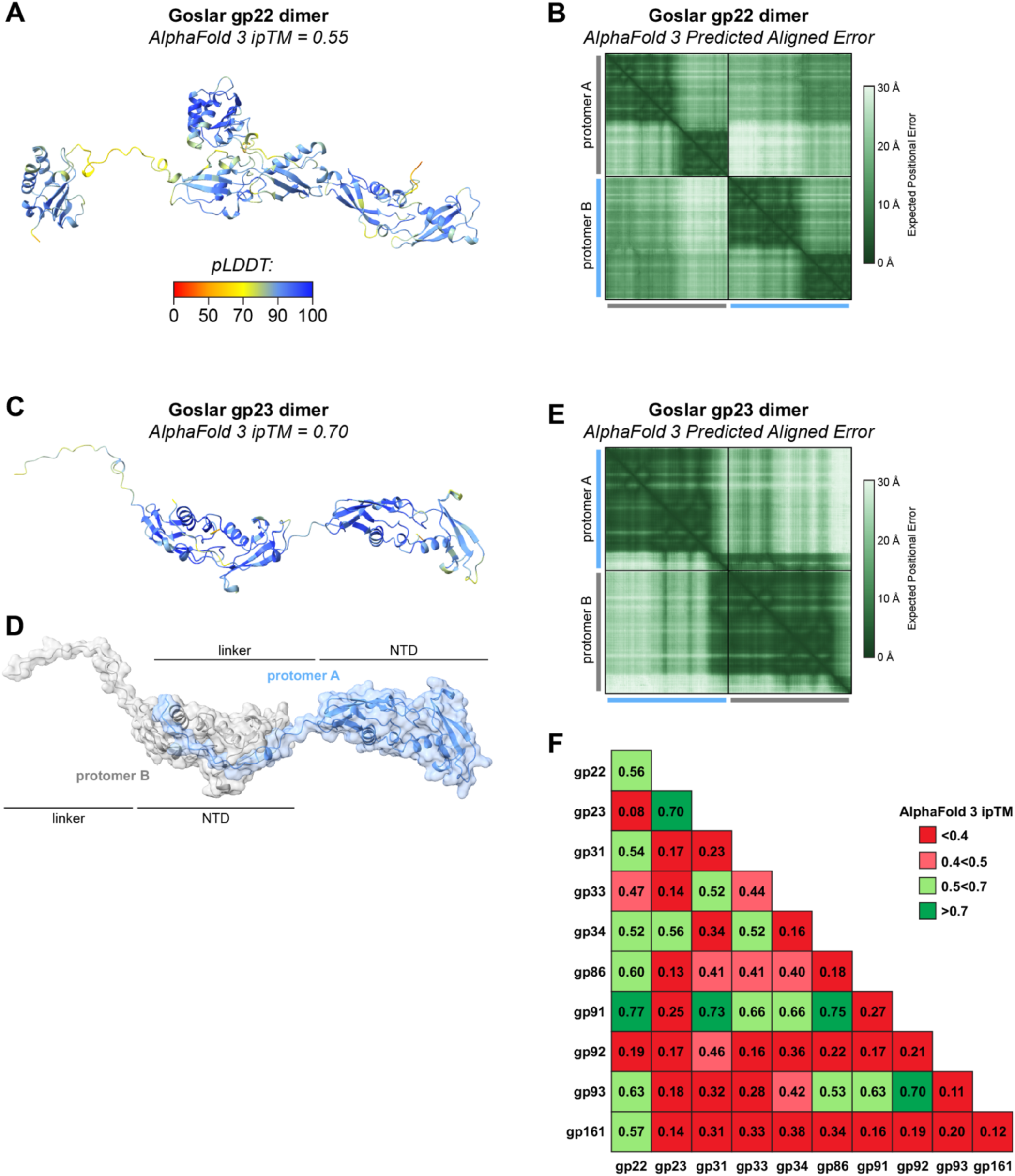
Oligomerization of DUF7941 proteins. A. AlphaFold 3 prediction of a Goslar gp22 dimer, colored by local confidence (pLDDT). View is equivalent to Figure 8B. B. AlphaFold 3 predicted aligned error (PAE) plot for the prediction of a Goslar gp22 dimer. C. AlphaFold 3 prediction of a Goslar gp23 dimer, colored by local confidence (pLDDT). D. AlphaFold 3 prediction of a Goslar gp23 dimer, colored by protomer (blue/gray) and with DUF7941 subdomains of each protomer indicated. E. AlphaFold 3 predicted aligned error (PAE) plot for the prediction of a Goslar gp23 dimer. F. AlphaFold 3 ipTM values for pairs of Goslar DUF7941 proteins. Shades of red indicate low confidence of interaction, and shades of red indicate increasing confidence of interaction.

**Figure S17.**
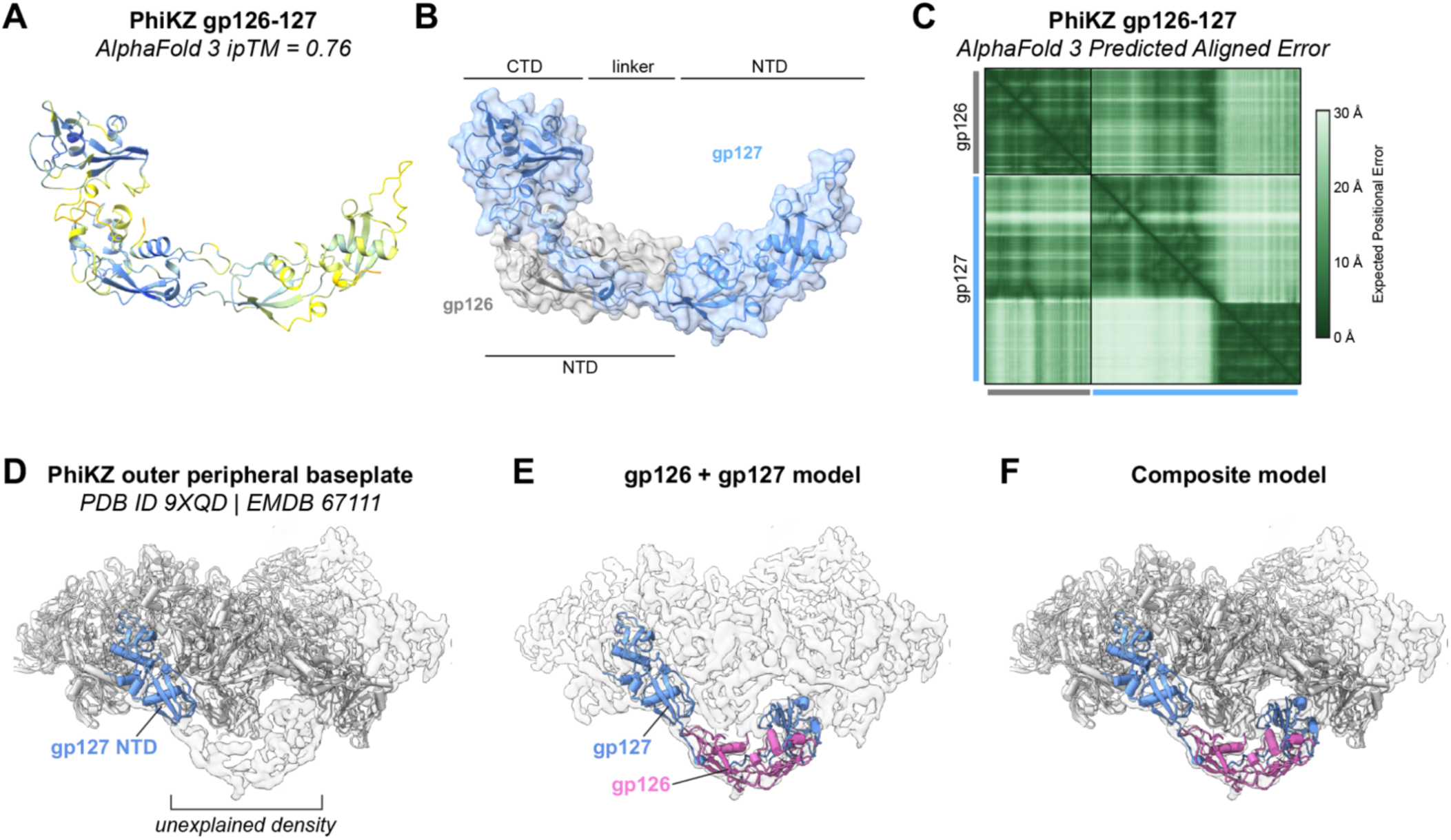
PhiKZ gp126-gp127 structure. A. AlphaFold 3 prediction of a PhiKZ gp126-gp127 heterodimer, colored by local confidence (pLDDT). B. AlphaFold 3 prediction of a PhiKZ gp126-gp127 heterodimer, colored by protein (gp127 blue, gp126 gray) and with DUF7941 subdomains of each protein indicated. C. AlphaFold 3 predicted aligned error (PAE) plot for the prediction of a PhiKZ gp126-gp127 heterodimer. D. PhiKZ outer peripheral baseplate structure (PDB ID 9QXD; EMDB-67111)^34^, with gp127 (NTD; residues 1-150) shown in blue. Unexplained density adjacent to the gp127 NTD model is noted. E. AlphaFold 3 model of the PhiKZ gp126-gp127 heterodimer manually fit into the outer peripheral baseplate map. Gp126 (pink) and the linker+CTD of gp127 (residues 151-290) fit into unexplained density. Fitting involved (1) overlaying the heterodimer model on the already-built gp127 NTD, (2) splitting the model between gp127 residues 150 and 151, then (3) rotating the (gp126 plus gp127 linker+CTD) coordinates ∼20° to fit the unexplained density. F. Composite model showing the PhiKZ outer peripheral baseplate model (gray) plus the gp126-gp127 heterodimer.

**Figure S18.**
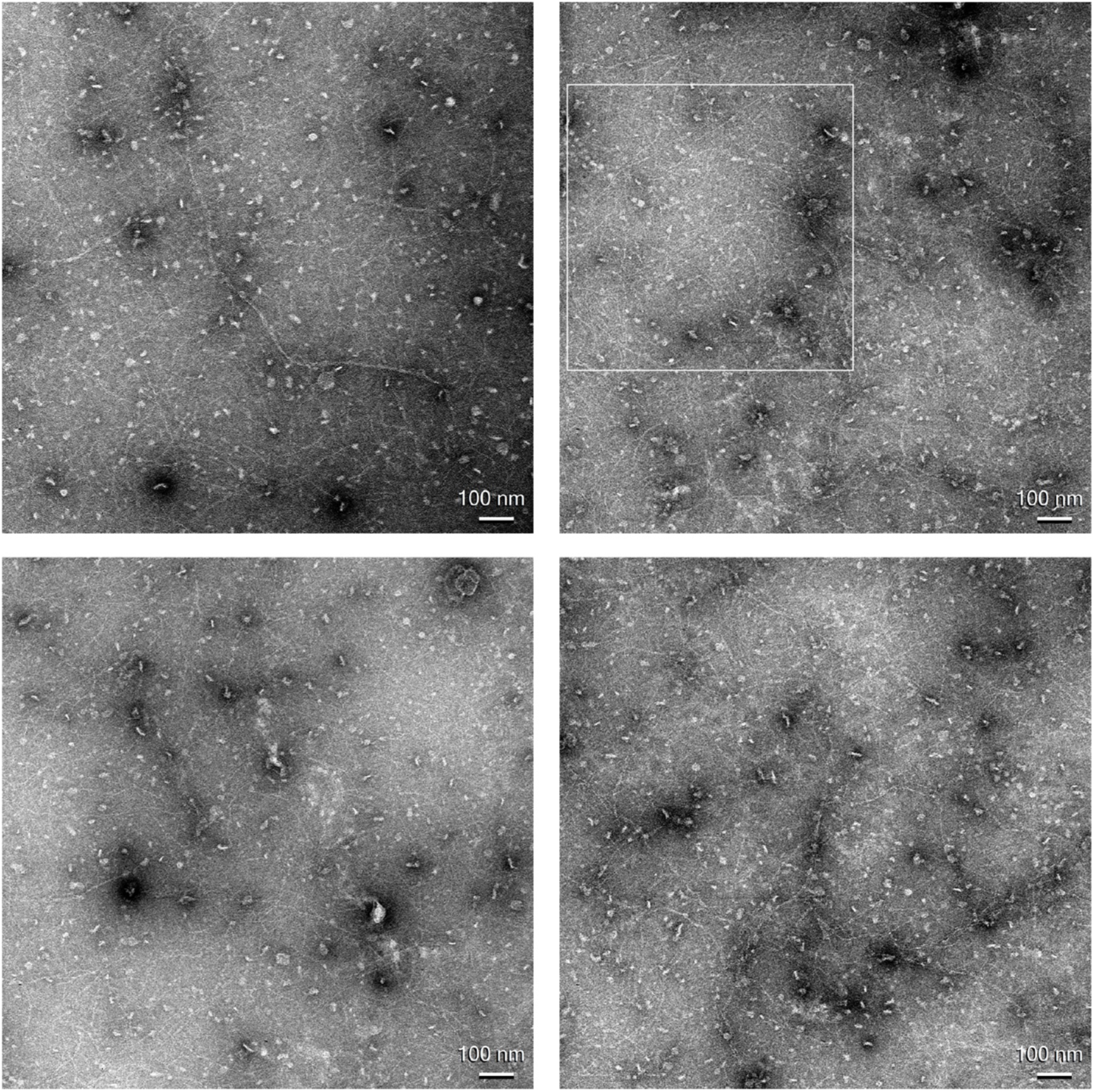
Negative stain-electron microscopy of Goslar gp22. Representative negative-stain electron microscopy images of purified Goslar gp22, showing filament formation. The white box on the upper-right image indicates the area shown in Figure 8C. Scale bar = 100 nm.

## REFERENCES

1. Keen, E.C. (2015). A century of phage research: bacteriophages and the shaping of modern biology. Bioessays 37, 6–9.

2. Hendrix, R.W. (2003). Bacteriophage genomics. Curr. Opin. Microbiol. 6, 506–511.

3. Doore, S.M., and Fane, B.A. (2016). The microviridae: Diversity, assembly, and experimental evolution. Virology 491, 45–55.

4. Al-Shayeb, B., Sachdeva, R., Chen, L.-X., Ward, F., Munk, P., Devoto, A., Castelle, C.J., Olm, M.R., Bouma-Gregson, K., Amano, Y., et al. (2020). Clades of huge phages from across Earth’s ecosystems. Nature 578, 425–431.

5. Chen, L. (2025). Discovery and analysis of an 841 kbp phage genome: the largest known to date. bioRxiv. 10.1101/2025.01.14.633092.

6. Crisci, M.A., Chen, L.-X., Devoto, A.E., Borges, A.L., Bordin, N., Sachdeva, R., Tett, A., Sharrar, A.M., Segata, N., Debenedetti, F., et al. (2021). Closely related Lak megaphages replicate in the microbiomes of diverse animals. iScience 24, 102875.

7. Saad, A.M., Soliman, A.M., Kawasaki, T., Fujie, M., Nariya, H., Shimamoto, T., and Yamada, T. (2019). Systemic method to isolate large bacteriophages for use in biocontrol of a wide-range of pathogenic bacteria. J. Biosci. Bioeng. 127, 73–78.

8. Kanhirun, N., Blanc, A., Aindow, A., You, M., Antani, J.D., Ghatbale, P., Leonard, J., Garcia, A.G., Nghiem, K., Whiteson, K., et al. (2024). Identification of a large cohort of Enterobacter jumbo phages with broad host ranges across pathogenic Gammaproteobacteria. bioRxiv. 10.1101/2024.10.29.620934.

9. M Iyer, L., Anantharaman, V., Krishnan, A., Burroughs, A.M., and Aravind, L. (2021). Jumbo phages: A comparative genomic overview of core functions and adaptions for biological conflicts. Viruses 13, 63.

10. Guan, J., and Bondy-Denomy, J. (2020). Intracellular organization by jumbo bacteriophages. J. Bacteriol. 203, e00362–20.

11. Chaikeeratisak, V., Nguyen, K., Khanna, K., Brilot, A.F., Erb, M.L., Coker, J.K.C., Vavilina, A., Newton, G.L., Buschauer, R., Pogliano, K., et al. (2017). Assembly of a nucleus-like structure during viral replication in bacteria. Science 355, 194–197.

12. Armbruster, E.G., Rani, P., Lee, J., Klusch, N., Hutchings, J., Hoffman, L.Y., Buschkaemper, H., Enustun, E., Adler, B.A., Inlow, K., et al. (2025). Sequential membrane- and protein-bound organelles compartmentalize genomes during phage infection. Cell Host Microbe 33, 484–497.e6.

13. Mozumdar, D., Fossati, A., Stevenson, E., Guan, J., Nieweglowska, E., Rao, S., Agard, D., Swaney, D.L., and Bondy-Denomy, J. (2024). Characterization of a lipid-based jumbo phage compartment as a hub for early phage infection. Cell Host Microbe 32, 1050–1058.e7.

14. Mendoza, S.D., Nieweglowska, E.S., Govindarajan, S., Leon, L.M., Berry, J.D., Tiwari, A., Chaikeeratisak, V., Pogliano, J., Agard, D.A., and Bondy-Denomy, J. (2020). A bacteriophage nucleus-like compartment shields DNA from CRISPR nucleases. Nature 577, 244–248.

15. Malone, L.M., Warring, S.L., Jackson, S.A., Warnecke, C., Gardner, P.P., Gumy, L.F., and Fineran, P.C. (2020). A jumbo phage that forms a nucleus-like structure evades CRISPR-Cas DNA targeting but is vulnerable to type III RNA-based immunity. Nat. Microbiol. 5, 48–55.

16. Antonova, D., Nichiporenko, A., Sobinina, M., Wang, Y., Vishnyakov, I.E., Moiseenko, A., Kurdyumova, I., Chesnokov, Y.M., Stepanchikova, E., Bourkaltseva, M., et al. (2024). Genomic transfer via membrane vesicle: a strategy of giant phage phiKZ for early infection. J. Virol. 98, e0020524.

17. Ceyssens, P.-J., Minakhin, L., Van den Bossche, A., Yakunina, M., Klimuk, E., Blasdel, B., De Smet, J., Noben, J.-P., Bläsi, U., Severinov, K., et al. (2014). Development of giant bacteriophage ϕKZ is independent of the host transcription apparatus. J. Virol. 88, 10501–10510.

18. Thomas, J.A., Rolando, M.R., Carroll, C.A., Shen, P.S., Belnap, D.M., Weintraub, S.T., Serwer, P., and Hardies, S.C. (2008). Characterization of Pseudomonas chlororaphis myovirus 201varphi2-1 via genomic sequencing, mass spectrometry, and electron microscopy. Virology 376, 330–338.

19. Yakunina, M., Artamonova, T., Borukhov, S., Makarova, K.S., Severinov, K., and Minakhin, L. (2015). A non-canonical multisubunit RNA polymerase encoded by a giant bacteriophage. Nucleic Acids Res. 43, 10411–10420.

20. Laughlin, T.G., Deep, A., Prichard, A.M., Seitz, C., Gu, Y., Enustun, E., Suslov, S., Khanna, K., Birkholz, E.A., Armbruster, E., et al. (2022). Architecture and self-assembly of the jumbo bacteriophage nuclear shell. Nature 608, 429–435.

21. Nieweglowska, E.S., Brilot, A.F., Méndez-Moran, M., Kokontis, C., Baek, M., Li, J., Cheng, Y., Baker, D., Bondy-Denomy, J., and Agard, D.A. (2023). The ϕPA3 phage nucleus is enclosed by a self-assembling 2D crystalline lattice. Nat. Commun. 14, 927.

22. Antonova, D., Belousova, V.V., Zhivkoplias, E., Sobinina, M., Artamonova, T., Vishnyakov, I.E., Kurdyumova, I., Arseniev, A., Morozova, N., Severinov, K., et al. (2023). The dynamics of synthesis and localization of jumbo phage RNA polymerases inside infected cells. Viruses 15, 2096.

23. Clark, S., Losick, R., and Pero, J. (1974). New RNA polymerase from Bacillus subtilis infected with phage PBS2. Nature 252, 21–24.

24. Sokolova, M., Borukhov, S., Lavysh, D., Artamonova, T., Khodorkovskii, M., and Severinov, K. (2017). A non-canonical multisubunit RNA polymerase encoded by the AR9 phage recognizes the template strand of its uracil-containing promoters. Nucleic Acids Res. 45, 5958–5967.

25. Prichard, A., Lee, J., Laughlin, T.G., Lee, A., Thomas, K.P., Sy, A., Spencer, T., Asavavimol, A., Cafferata, A., Cameron, M., et al. (2023). Identifying the core genome of the nucleus-forming bacteriophage family and characterization of Erwinia phage RAY. Cell Rep. 42, 112432.

26. Chaikeeratisak, V., Khanna, K., Nguyen, K.T., Egan, M.E., Enustun, E., Armbruster, E., Lee, J., Pogliano, K., Villa, E., and Pogliano, J. (2022). Subcellular organization of viral particles during maturation of nucleus-forming jumbo phage. Sci. Adv. 8, eabj9670.

27. Prichard, A., and Pogliano, J. (2024). The intricate organizational strategy of nucleus-forming phages. Curr. Opin. Microbiol. 79, 102457.

28. Birkholz, E.A., Laughlin, T.G., Armbruster, E., Suslov, S., Lee, J., Wittmann, J., Corbett, K.D., Villa, E., and Pogliano, J. (2022). A cytoskeletal vortex drives phage nucleus rotation during jumbo phage replication in E. coli. Cell Rep. 40, 111179.

29. Enustun, E., Armbruster, E.G., Lee, J., Zhang, S., Yee, B.A., Malukhina, K., Gu, Y., Deep, A., Naritomi, J.T., Liang, Q., et al. (2024). A phage nucleus-associated RNA-binding protein is required for jumbo phage infection. Nucleic Acids Res. 52, 4440–4455.

30. Morgan, C.J., Enustun, E., Armbruster, E.G., Birkholz, E.A., Prichard, A., Forman, T., Aindow, A., Wannasrichan, W., Peters, S., Inlow, K., et al. (2024). An essential and highly selective protein import pathway encoded by nucleus-forming phage. Proc. Natl. Acad. Sci. U. S. A. 121, e2321190121.

31. Kokontis, C., Klein, T.A., Silas, S., and Bondy-Denomy, J. (2025). Multi-interface licensing of protein import into a phage nucleus. Nature 639, 456–462.

32. Fokine, A., Battisti, A.J., Bowman, V.D., Efimov, A.V., Kurochkina, L.P., Chipman, P.R., Mesyanzhinov, V.V., and Rossmann, M.G. (2007). Cryo-EM study of the Pseudomonas bacteriophage phiKZ. Structure 15, 1099–1104.

33. Chan, B.K., Stanley, G.L., Kortright, K.E., Vill, A.C., Modak, M., Ott, I.M., Sun, Y., Würstle, S., Grun, C.N., Kazmierczak, B.I., et al. (2025). Personalized inhaled bacteriophage therapy for treatment of multidrug-resistant Pseudomonas aeruginosa in cystic fibrosis. Nat. Med. 31, 1494–1501.

34. Xiao, H., Peng, Z., Zhou, J., Chen, Y., Peng, Y., Tang, Y., Li, T., Chen, W., Huang, S.-Y., Cheng, L., et al. (2026). Structural atlas of the intact jumbo phage phiKZ. Nat. Commun. 10.1038/s41467-026-71561-2.

35. Punjani, A., Rubinstein, J.L., Fleet, D.J., and Brubaker, M.A. (2017). cryoSPARC: algorithms for rapid unsupervised cryo-EM structure determination. Nat. Methods 14, 290–296.

36. Jamali, K., Käll, L., Zhang, R., Brown, A., Kimanius, D., and Scheres, S.H.W. (2024). Automated model building and protein identification in cryo-EM maps. Nature 628, 450–457.

37. Abramson, J., Adler, J., Dunger, J., Evans, R., Green, T., Pritzel, A., Ronneberger, O., Willmore, L., Ballard, A.J., Bambrick, J., et al. (2024). Accurate structure prediction of biomolecular interactions with AlphaFold 3. Nature 630, 493–500.

38. Wu, W., Thomas, J.A., Cheng, N., Black, L.W., and Steven, A.C. (2012). Bubblegrams reveal the inner body of bacteriophage φKZ. Science 335, 182.

39. Krylov, V.N., Smirnova, T.A., Minenkova, I.B., Plotnikova, T.G., Zhazikov, I.Z., and Khrenova, E.A. (1984). Pseudomonas bacteriophage phi KZ contains an inner body in its capsid. Can. J. Microbiol. 30, 758–762.

40. Thomas, J.A., Weintraub, S.T., Wu, W., Winkler, D.C., Cheng, N., Steven, A.C., and Black, L.W. (2012). Extensive proteolysis of head and inner body proteins by a morphogenetic protease in the giant Pseudomonas aeruginosa phage φKZ: Extensive head protein proteolysis in myovirus φKZ. Mol. Microbiol. 84, 324–339.

41. Yang, Y., Shao, Q., Guo, M., Han, L., Zhao, X., Wang, A., Li, X., Wang, B., Pan, J.-A., Chen, Z., et al. (2024). Capsid structure of bacteriophage ΦKZ provides insights into assembly and stabilization of jumbo phages. Nat. Commun. 15, 6551.

42. Nichiporenko, A., Antonova, D., Kurdyumova, I., Khodorkovskii, M., and Yakunina, M.V. (2024). Assembly of phiKZ bacteriophage Inner Body during infection. Biochem. Biophys. Res. Commun. 693, 149372.

43. Duda, R.L., and Teschke, C.M. (2019). The amazing HK97 fold: versatile results of modest differences. Curr. Opin. Virol. 36, 9–16.

44. Helgstrand, C., Wikoff, W.R., Duda, R.L., Hendrix, R.W., Johnson, J.E., and Liljas, L. (2003). The refined structure of a protein catenane: The HK97 bacteriophage capsid at 3.44 Å resolution. J. Mol. Biol. 334, 885–899.

45. Evseev, P., Shneider, M., and Miroshnikov, K. (2022). Evolution of phage tail sheath protein. Viruses 14, 1148.

46. Zinke, M., Schröder, G.F., and Lange, A. (2022). Major tail proteins of bacteriophages of the order Caudovirales. J. Biol. Chem. 298, 101472.

47. Taylor, N.M.I., Prokhorov, N.S., Guerrero-Ferreira, R.C., Shneider, M.M., Browning, C., Goldie, K.N., Stahlberg, H., and Leiman, P.G. (2016). Structure of the T4 baseplate and its function in triggering sheath contraction. Nature 533, 346–352.

48. Guan, J., Oromí-Bosch, A., Mendoza, S.D., Karambelkar, S., Berry, J.D., and Bondy-Denomy, J. (2022). Bacteriophage genome engineering with CRISPR-Cas13a. Nat. Microbiol. 7, 1956–1966.

49. Holm, L. (2022). Dali server: structural unification of protein families. Nucleic Acids Res. 50, W210–W215.

50. van Kempen, M., Kim, S.S., Tumescheit, C., Mirdita, M., Lee, J., Gilchrist, C.L.M., Söding, J., and Steinegger, M. (2024). Fast and accurate protein structure search with Foldseek. Nat. Biotechnol. 42, 243–246.

51. Jha, N., Kravitz, J., West-Roberts, J., Lu, C., Camargo, A.P., Roux, S., Cornman, A., and Hwang, Y. (2025). Gaia: An AI-enabled genomic context-aware platform for protein sequence annotation. Sci. Adv. 11, eadv5109.

52. Conley, M.P., and Wood, W.B. (1975). Bacteriophage T4 whiskers: a rudimentary environment-sensing device. Proc. Natl. Acad. Sci. U. S. A. 72, 3701–3705.

53. Fokine, A., Zhang, Z., Kanamaru, S., Bowman, V.D., Aksyuk, A.A., Arisaka, F., Rao, V.B., and Rossmann, M.G. (2013). The molecular architecture of the bacteriophage T4 neck. J. Mol. Biol. 425, 1731–1744.

54. Noteborn, W.E.M., Ouyang, R., Hoeksma, T., Sidi Mabrouk, A., Esteves, N.C., Pelt, D.M., Scharf, B.E., and Briegel, A. (2025). Insights into the structure and initial host attachment of the flagellotropic bacteriophage 7-7-1. Commun. Biol. 9, 55.

55. Erb, M.L., Kraemer, J.A., Coker, J.K.C., Chaikeeratisak, V., Nonejuie, P., Agard, D.A., and Pogliano, J. (2014). A bacteriophage tubulin harnesses dynamic instability to center DNA in infected cells. Elife 3. 10.7554/eLife.03197.

56. Harding, K.R., Malone, L.M., Kyte, N.A.P., Jackson, S.A., Smith, L.M., and Fineran, P.C. (2025). Genome-wide identification of bacterial genes contributing to nucleus-forming jumbo phage infection. Nucleic Acids Res. 53. 10.1093/nar/gkae1194.

57. Ranta, K., Skurnik, M., and Kiljunen, S. (2025). Isolation and characterization of fMGyn-Pae01, a phiKZ-like jumbo phage infecting Pseudomonas aeruginosa. Virol. J. 22, 55.

58. Emsley, P., Lohkamp, B., Scott, W.G., and Cowtan, K. (2010). Features and development of Coot. Acta Crystallogr. D Biol. Crystallogr. 66, 486–501.

59. Afonine, P.V., Poon, B.K., Read, R.J., Sobolev, O.V., Terwilliger, T.C., Urzhumtsev, A., and Adams, P.D. (2018). Real-space refinement in PHENIX for cryo-EM and crystallography. Acta Crystallogr. D Struct. Biol. 74, 531–544.

60. Meng, E.C., Goddard, T.D., Pettersen, E.F., Couch, G.S., Pearson, Z.J., Morris, J.H., and Ferrin, T.E. (2023). UCSF ChimeraX: Tools for structure building and analysis. Protein Sci. 32, e4792.

61. Blum, M., Andreeva, A., Florentino, L.C., Chuguransky, S.R., Grego, T., Hobbs, E., Pinto, B.L., Orr, A., Paysan-Lafosse, T., Ponamareva, I., et al. (2025). InterPro: the protein sequence classification resource in 2025. Nucleic Acids Res. 53, D444–D456.

62. Elfmann, C., and Stülke, J. (2023). PAE viewer: a webserver for the interactive visualization of the predicted aligned error for multimer structure predictions and crosslinks. Nucleic Acids Res. 51, W404–W410.

63. Zhang, J., Xin, L., Shan, B., Chen, W., Xie, M., Yuen, D., Zhang, W., Zhang, Z., Lajoie, G.A., and Ma, B. (2012). PEAKS DB: de novo sequencing assisted database search for sensitive and accurate peptide identification. Mol. Cell. Proteomics 11, M111.010587.

